# A role for glutathione in buffering excess intracellular copper in *Streptococcus pyogenes*

**DOI:** 10.1101/2020.05.14.095349

**Authors:** Louisa J. Stewart, Cheryl-lynn Y. Ong, May M. Zhang, Stephan Brouwer, Liam McIntyre, Mark R. Davies, Mark J. Walker, Alastair G. McEwan, Kevin J. Waldron, Karrera Y. Djoko

**Affiliations:** Department of Biosciences, Durham University, Durham DH1 3LE, United Kingdom; School of Chemistry and Molecular Biosciences and Australian Infectious Diseases Research Centre, The University of Queensland, St Lucia, QLD 4072, Australia; Department of Microbiology and Immunology, University of Melbourne, at the Peter Doherty Institute for Infection and Immunity, Melbourne, VIC 3000, Australia; Biosciences Institute, Faculty of Medical Sciences, Framlington Place, Newcastle University, Newcastle upon Tyne, NE2 4HH, United Kingdom

**Author notes:** Corresponding author Mailing address: Department of Biosciences, Durham University, Durham DH1 3LE, United Kingdom. We consider that these authors have contributed equally to the work (Louisa J. Stewart, Cheryl-lynn Y. Ong).

**Keywords:** copper homeostasis, copper stress and tolerance, copper export, metal buffer, glutathione, Group A *Streptococcus*

## Abstract

Copper (Cu) is an essential metal for bacterial physiology but in excess it is bacteriotoxic. To limit Cu levels in the cytoplasm, most bacteria possess a transcriptionally-responsive system for Cu export. In the Gram-positive human pathogen *Streptococcus pyogenes* (Group A *Streptococcus*, GAS), this system is encoded by the *copYAZ* operon. In this study, we demonstrate that the site of GAS infection *in vivo* represents a Cu-rich environment but inactivation of the *copA* Cu efflux gene does not reduce virulence in a mouse model of invasive disease. *In vitro*, Cu treatment leads to multiple observable phenotypes, including defects in growth and viability, decreased fermentation, inhibition of glyceraldehyde 3-phosphate dehydrogenase (GapA) activity, and misregulation of metal homeostasis, likely as a consequence of mismetalation of non-cognate metal-binding sites. Surprisingly, the onset of these effects is delayed by ∼4 h even though expression of *copZ* is upregulated immediately upon exposure to Cu. We further show that the onset of all phenotypes coincides with depletion of intracellular glutathione (GSH). Supplementation with extracellular GSH replenishes the intracellular pool of this thiol and suppresses all the observable effects of Cu treatment. Our results indicate that GSH contributes to buffering of excess intracellular Cu when the transcriptionally-responsive Cu export system is overwhelmed. Thus, while the *copYAZ* operon is responsible for Cu *homeostasis*, GSH has a role in Cu *tolerance* that allows bacteria to maintain metabolism even in the presence of an excess of this metal ion. This study advances fundamental understanding of Cu handling in the bacterial cytoplasm.

**IMPORTANCE:** The control of intracellular metal availability is fundamental to bacterial physiology. In the case of copper (Cu), it is established that rising intracellular Cu levels eventually fill the metal-sensing site of the endogenous Cu-sensing transcriptional regulator, which in turn induces transcription of a copper export pump. This response caps intracellular Cu availability below a well-defined threshold and prevents Cu toxicity. Glutathione, abundant in many bacteria, is known to bind Cu and is long assumed to contribute to bacterial Cu handling. However, there is some ambiguity since neither its biosynthesis nor uptake is Cu-regulated. Furthermore, there is little experimental support for this role of glutathione beyond measurement of the effect of Cu on growth of glutathione-deficient mutants. Our work with Group A *Streptococcus* provides new evidence that glutathione increases the threshold of intracellular Cu availability that can be tolerated by bacteria and thus advances fundamental understanding of bacterial Cu handling.

## INTRODUCTION

Bacteria have been exposed to environmental copper (Cu) since the Great Oxidation Event, when the rise in atmospheric O_2_ levels led to solubilisation of Cu from minerals. There is evidence that recent evolution of plant, animal, and human pathogens has been influenced by the anthropogenic release of Cu into soils, for instance *via* mining activities and the legacy of using Cu salts and compounds in industrial-scale biocides (1). Bacteria also encounter elevated levels of Cu at microenvironments within a eukaryotic host. Bacterial predation induces an increase in intracellular Cu levels in protozoa (2) while phagocytosis stimulates uptake and accumulation of Cu in murine macrophages (3, 4). Studies of more complex animal models of infectious disease and human infections further suggest that infection triggers systemic changes in host Cu levels and that the specific sites of inflammation are usually, though not always, Cu-rich (5–9). The prevailing model that encompasses these observations suggests that Cu exerts a direct antibacterial action and/or supports the antibacterial function of innate immune cells (10).

Cu can be bacteriotoxic because it is a competitive metal for protein binding (11). Extracellular Cu invariably enters the bacterial cytoplasm *via* an uptake process that remains poorly understood. Once inside, Cu fills the available Cu-binding sites in proteins and other biomolecules, beginning with the tightest affinity and eventually associating with the weakest affinity sites. Within this hierarchy of binding sites are the allosteric sites in Cu-sensing transcriptional regulators, which, when metalated by Cu, activate expression of a Cu efflux pump (12). In undertaking this role, the Cu sensor and export pump together impose an upper threshold of Cu availability in the cytoplasm. They ensure that only native, stable, high-affinity Cu sites are metalated by Cu, and at the same time they prevent adventitious, non-specific, non-cognate, weaker-binding sites from becoming mismetalated. Such mismetalation events can inactivate key enzymes and, consequently, impair bacterial growth and viability (13–16).

Additional cytoplasmic components are thought to limit Cu availability by chelating or “buffering” this metal ion. These components include bacterial metallothioneins (17), copper storage proteins (18), and Cu-binding metallochaperones (19, 20), which are often, though not always, transcriptionally regulated by the endogenous Cu sensors. Mutant bacterial strains lacking these proteins typically display a Cu-sensitive growth phenotype. Non-protein components, particularly the low molecular weight thiol glutathione (GSH), are also assumed to buffer Cu (21), although their uptake or biosynthesis is not transcriptionally induced in response to Cu treatment (15, 20, 22, 23). *In vitro*, addition of GSH protects purified metalloenzymes from inactivation by Cu (13). *In vivo*, growth of bacterial mutant strains that are impaired in GSH uptake (24) or biosynthesis (25–27) are all inhibited by added Cu, especially if the Cu efflux pump (25, 26) or Cu-binding metallochaperone (27) in the organism is also inactivated.

Beyond growth analysis, there is currently little experimental support for a role of GSH in buffering Cu in bacteria. Perhaps the clearest, albeit indirect, line of evidence was obtained using a Δ*gshB*Δ*atx1* mutant of *Synechocystis* lacking the GSH biosynthesis enzyme GshB and the cytoplasmic Cu-binding metallochaperone Atx1. This mutant failed to repress expression of Zn-regulated genes in response to elevated Zn (27). *In vitro* metal- and DNA-binding experiments (28) suggest that the absence of GSH and the metallochaperone leads to an increase in background Cu availability, which mismetalates the allosteric site of the Zn sensor Zur and thus interferes with Zn sensing.

Like most bacteria, Gram-positive human pathogen *Streptococcus pyogenes* (Group A *Streptococcus*, GAS) possesses a system for Cu sensing and efflux, which is encoded by the *copYAZ* operon (29). In this work, we examine whether *copA*, encoding the Cu-effluxing P_1B-1_-type ATPase, plays a critical role in GAS pathogenesis, as demonstrated for other bacterial pathogens (7, 30–32). We show that GAS occupies a Cu-rich environment during infection of a mouse model of invasive disease, and yet inactivation of *copA* does not significantly reduce GAS virulence. This unexpected observation leads us to investigate the effects of Cu treatment on the cellular biochemistry and physiology of GAS. The results provide key insights into the importance of GSH in cytoplasmic Cu buffering to supplement the transcriptionally-responsive Cu sensing and efflux system. This additional buffering extends the range of intracellular Cu concentrations that can be tolerated by bacteria, and thus prevents a sudden or abrupt transition from Cu homeostasis to Cu stress upon exposure to an excess of this metal ion.

## RESULTS

### Initial characterisation of a Δ*copA* mutant

The *copYAZ* operon in GAS has been previously shown to resemble other Cop systems in Gram-positive bacteria (29) (Supporting Figure 1A). Consistent with a role in Cu efflux, expression of this operon functionally complemented a heterologous *Escherichia coli* Δ*copA* mutant strain (29). Our *in silico* analyses found one additional open reading frame downstream of *copZ* (Supporting Figure 1A). It encodes a small, uncharacterised protein (56 amino acids) with an N-terminal transmembrane domain, a putative metal-binding C-X_3_-M-H motif at the C-terminus, and no known homologue. This gene is absent from *copYAZ* operons in other Gram-positive bacteria and its function in Cu homeostasis is unknown.

For the present study, we constructed a non-polar Δ*copA* mutant of GAS M1T1 strain 5448. This mutation did not alter basal expression of downstream *cop* genes (Supporting Figure 1B(i)). As anticipated, the Δ*copA* mutant was more susceptible to growth inhibition by added Cu than was the wild type (Supporting Figure 1C). This mutant also accumulated higher levels of intracellular Cu (Supporting Figure 1D), leading to increased expression of the other *cop* genes when compared with the wild type (Supporting Figure 1B(ii)). Marker rescue (*copA*^+^) restored both the expression of *copA* and the wild type phenotype (Supporting Figures 1B-D).

### Deletion of *copA* does not lead to a loss of virulence in a mouse model of infection

To determine if the Cop system and interactions with host Cu have an effect on GAS pathogenesis, we employed an established invasive disease model using transgenic human-plasminogenised mice (33). Mice subcutaneously infected with wild type GAS developed ulcerative skin lesions at the site of injection after 1 day. These lesions were excised 3 days post-infection and were found to contain more Cu than adjacent healthy skin or skin from uninfected mice (Figure 1A). Consistent with these results, the *copYAZ* operon was upregulated in GAS isolated from infected mouse tissues when compared with those grown in THY medium (34). There was also an increase in Cu levels in mouse blood after 3 days of infection (Figure 1B). Notably, these Cu levels in the blood are comparable to those measured in the serum of mice infected with the fungal pathogen *Candida albicans* or the parasite *Plasmodium berghei* (5). These observations support a model where redistribution of host Cu is a feature of the general immune response to infection (5).

**Figure 1.**
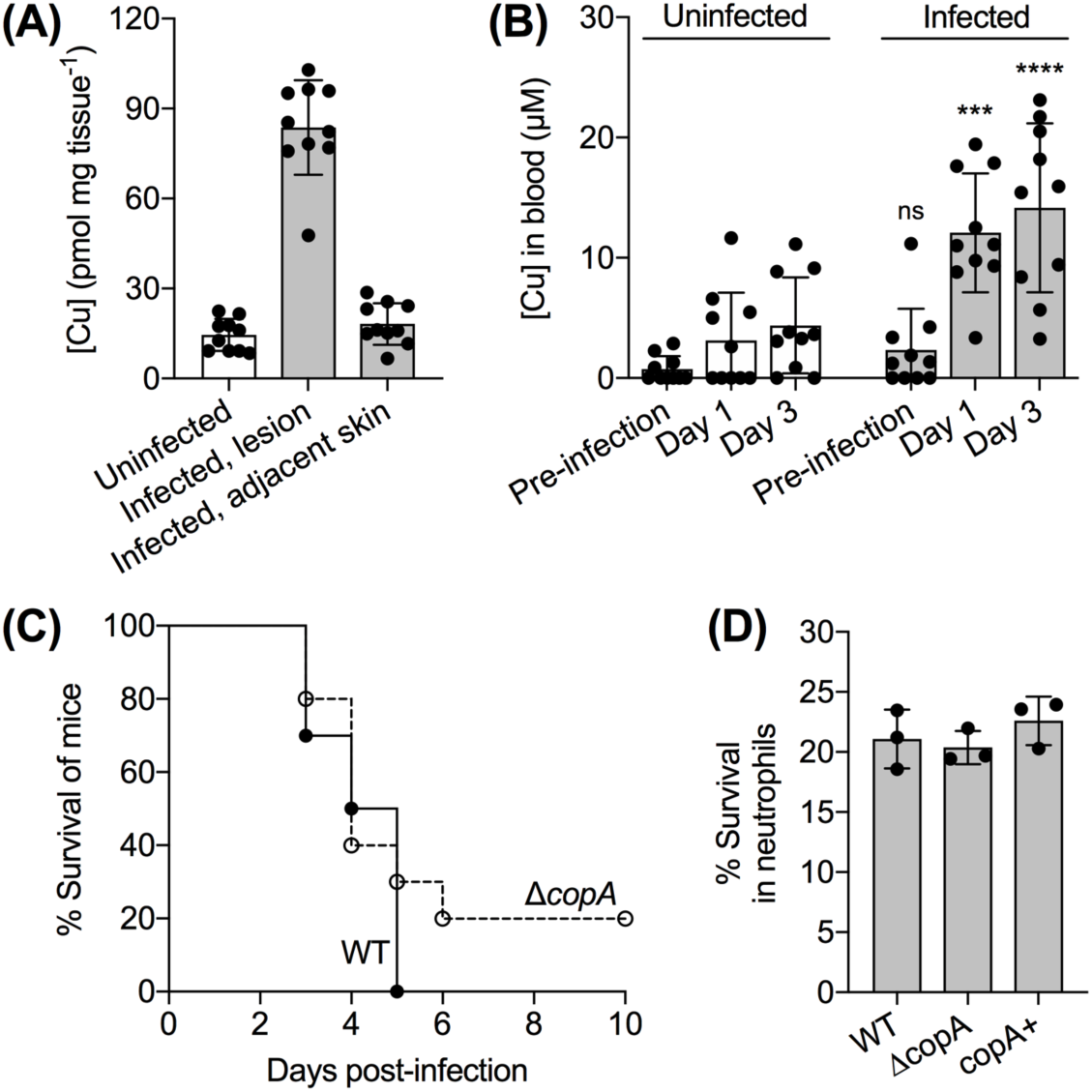
Changes in Cu levels during GAS infection and the effect of a *copA* mutation on GAS virulence in host infection models. **(A) Cu levels in mouse lesions.** Mice were infected subcutaneously with GAS wild type strain or left uninfected (*n* = 10 each). After 3 days, skin from uninfected mice, and both skin lesions and healthy skin adjacent to the lesions were excised. Total Cu levels were measured by ICP MS and normalised to the weight of the tissues. Cu levels in infected lesions were higher than those in adjacent healthy skin (*P* < 0.0001) or skin from uninfected mice (*P* < 0.0001). **(B) Cu levels in mouse blood.** Mice were infected subcutaneously with GAS wild type strain or left uninfected (*n* = 10 each). Blood was collected and total Cu levels were measured by ICP MS. Values below the detection limit were represented as zero. Cu levels in the blood of infected mice on Days 1 and 3 were higher from those in the blood of uninfected mice (\*\*\**P* = 0.0001, \*\*\*\**P* < 0.0001). ^ns^*P* = 0.81 (*vs.* uninfected mice). **(C) Virulence in an *in vivo* mouse model of infection.** Mice were infected subcutaneously with GAS wild type (WT) or Δ*copA* mutant strains (*n* = 10 each). The number of surviving mice was counted daily up to 10 days post-infection. Differences in survival curves were analysed using the Mann-Whitney test, which found no statistical difference (*P* = 0.099). **(D) Virulence in an *ex vivo* human neutrophil model of infection.** Human neutrophils were infected with GAS wild type (WT), Δ*copA*, or *copA*^+^ mutant strains (*n* = 3 each). Survival of bacteria relative to the input was measured after 2 h. There was no difference between survival of the Δ*copA* mutant when compared with the WT (*P* = 0.87) or *copA*^+^ (*P* = 0.35) strains.

Comparing the survival of mice post-infection, no statistically significant difference was observed whether mice were infected with wild type or the *ΔcopA* mutant (*P* = 0.0991; Figure 1C). Although no single animal model can fully represent the complex features of human streptococcal diseases (35), consistent with *in vivo* findings, the Δ*copA* mutant was no more susceptible to killing by human neutrophils when compared with the wild type or *copA*^+^ mutant strains in an *ex vivo* infection assay (Figure 1D). In addition, recent reports did *not* identify the *cop* genes to be fitness determinants during *ex vivo* infection of human blood (36) or *in vivo* soft tissue infection in mice (37). These results imply that, despite the systemic and niche-specific elevated levels of host Cu, the Cu efflux pump CopA is not essential for GAS virulence in this model.

### Cu treatment leads to defects in the late exponential phase of growth

The lack of a virulence defect for the *ΔcopA* mutant *in vivo* prompted us to examine the impact of Cu treatment on GAS physiology *in vitro*. Addition of Cu (up to 10 µM) to the culture medium did not affect the doubling time of the Δ*copA* mutant during the exponential phase of growth but it did reduce the final culture yield (Figure 2A, Supporting Figures 2A-B). This phenotype was reproduced during growth in the presence of glucose or alternative carbon sources (Supporting Figure 2C). In each condition, growth of Cu-treated cultures ceased upon reaching approximately the same OD_600_ (∼0.35) regardless of growth rate, indicating that the growth defect was related to bacterial cell numbers and/or growth stage. Consistent with this interpretation, Cu treatment did not affect growth in the presence of mannose (Supporting Figures 2C) or limiting amounts of glucose (Supporting Figures 2D), since neither experimental condition supported growth of GAS beyond OD_600_ ∼0.35.

**Figure 2.**
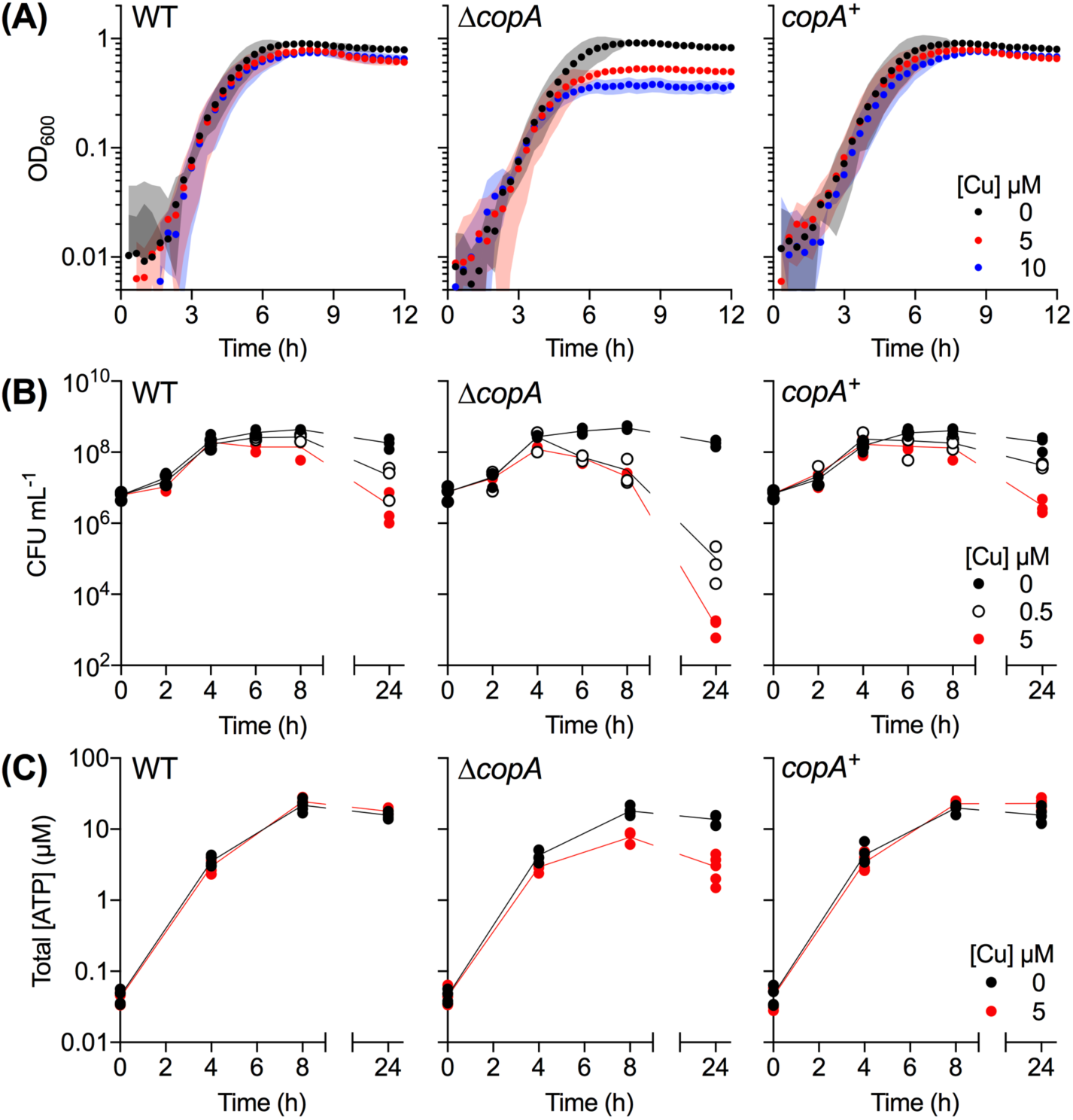
Cu-dependent defects in growth and viability. GAS strains were cultured with added Cu as indicated. **(A) Growth.** Cultures (*n* = 3) were grown in microtitre plates and OD_600_ values were recorded every 20 min. Cu treatment clearly suppressed growth of Δ*copA* cultures (*P* = 0.034 for 5 µM Cu, *P* < 0.0001 for 10 µM Cu). **(B) Plating efficiency.** Cultures (*n* = 3) were plated out at the indicated time points and the number of colony-forming units (CFU) was enumerated. Cu treatment clearly suppressed plating efficiency of the Δ*copA* cultures (*P* < 0.0001 for both 0.5 and 5 µM Cu). **(C) Total ATP levels.** Cultures (*n* = 5) were sampled at the indicated time points and total ATP levels were determined. Cu treatment clearly suppressed ATP production in the Δ*copA* cultures (*P* < 0.0001). All statistical analyses were *vs.* 0 µM Cu.

Parallel assessments of plating efficiency and total ATP levels confirmed that differences between Cu-treated and untreated cultures appeared only in the late exponential or early stationary phase of growth (after ∼4 h when grown in the presence of glucose; Figures 2B-C). There were clear decreases in the plating efficiency and ATP production by Cu-treated Δ*copA* cultures during this period when compared with the untreated control.

### Cu treatment leads to metabolic arrest in the late exponential phase of growth

GAS is a lactic acid bacterium. Under our experimental conditions, this organism carried out homolactic fermentation and generated lactic acid as the major end product (Supporting Figures 3A-B). However, we noted that Cu-treated Δ*copA* cultures did not acidify the growth medium (Supporting Figure 3C). Hence, we hypothesised that Cu treatment impairs fermentation in GAS.

Consistent with this proposal, Cu-treated Δ*copA* cultures produced ∼50% less lactic acid and consumed ∼50% less glucose when compared with the untreated control (Figure 3A, Supporting Figures 3B(i)-(ii)). Pyruvate production remained unaltered (Supporting Figure 3B(iii)). There is no evidence of a shift towards mixed-acid fermentation since the reduction in lactate levels was not accompanied by a concomitant increase in acetate levels (Supporting Figure 3B(iv)). The levels of ethanol were undetectable (detection limit ∼0.2 mM).

**Figure 3.**
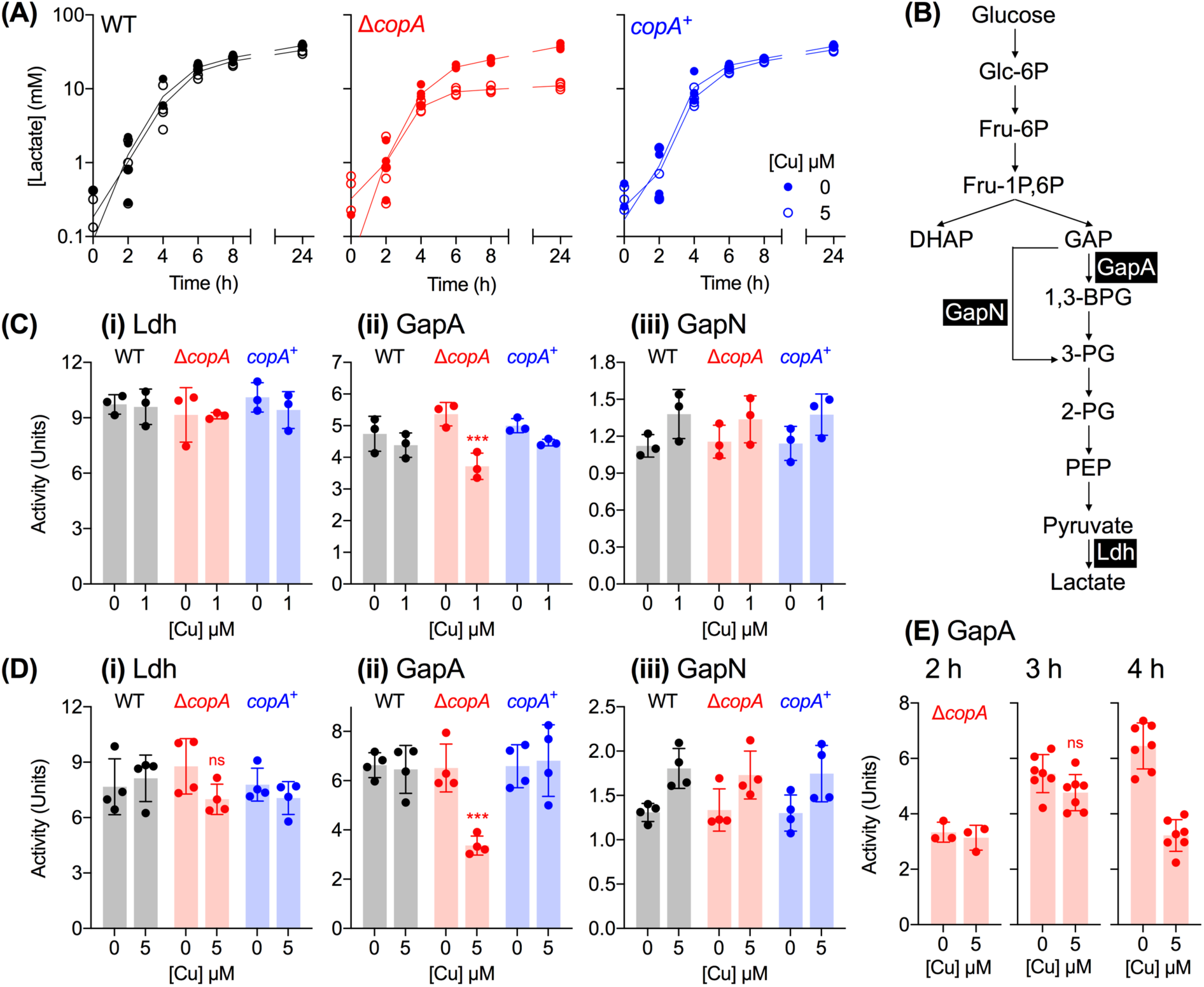
Cu-dependent defects in glycolysis and homolactic fermentation. **(A) Lactate production.** GAS strains were cultured with added Cu as indicated (*n* = 3). Amounts of lactate secreted to the extracellular culture medium were measured at the indicated time points. Cu treatment clearly suppressed lactate production in the Δ*copA* cultures (*P* < 0.0001). **(B) Fermentation pathway in GAS.** Enzymes of interest, namely GapA (NAD^+^-dependent GAPDH, M5005_SPy_0233), GapN (NADP^+^-dependent GAPDH, M5005_SPy_1119), and Ldh (lactate dehydrogenase, M5005_SPy_0873) are shown. **(C)-(D) Activity of glycolytic enzymes (i) Ldh**, **(ii) GapA**, **and (iii) GapN.** GAS strains were cultured for t = 4 h with **(C)** 0 or 1 µM of added Cu (*n* = 3), or **(D)** 0 or 5 µM of added Cu (*n* = 4). Enzyme activities were determined in cell extracts. Cu treatment clearly decreased GapA activity in Δ*copA* cultures (\*\*\**P* = 0.0004). ^ns^*P* = 0.14. **(E) GapA activity over time.** GAS Δ*copA* mutant strain was cultured with added Cu as indicated for t = 2 h (*n* = 3), 3 h (*n* = 7), or 4 h (*n* = 7). Enzyme activities were determined in cell extracts. Cu treatment did not have an effect on GapA activity at t = 2 h (*P* = 0.99) or 3 h (^ns^*P* = 0.18), but it strongly inhibited GapA activity at t = 4 h (*P* < 0.0001). All statistical analyses were *vs.* 0 µM Cu.

Differences in lactate production between Cu-treated and untreated Δ*copA* cultures appeared, again, only after ∼4 h of growth (Figure 3A). While our methods are not sufficiently sensitive to detect small changes in glucose levels at earlier time points, it is clear that Cu-treated Δ*copA* cultures did not consume glucose beyond t ∼ 4 h (Supporting Figure 3D(i)). Pyruvate production was, again, not affected at any time point (Supporting Figure 3D(ii)). These results suggest that Cu treatment leads to defects in metabolism but only after entry into the late exponential phase of growth.

### Cu treatment results in a loss in GapA activity in the late exponential phase of growth

The loss in lactate production, but not pyruvate, implies that lactate dehydrogenase (Ldh) is inactivated (Figure 3B). To test this proposal, we cultured GAS in the absence or presence of added Cu for 4 h, prepared whole cell extracts, and measured Ldh activity. Figures 3C(i) and 3D(i) show that Ldh remained active in all strains, regardless of Cu treatment (0, 1, or 5 µM of added Cu).

What, then, is the target of Cu intoxication in GAS? This bacterium does not possess a TCA cycle or the biosynthesis pathways for multiple amino acids, vitamins, and cofactors (e.g. heme). Thus, it lacks obvious candidate iron-sulfur cluster enzymes that are destabilised by excess Cu ions in other systems (13). In an attempt to develop a molecular explanation for the loss in fermentation, we examined the activity of the two GAPDH enzymes in GAS, namely the classical, phosphorylating, ATP-generating GapA and the alternative, non-phosphorylating GapN (Figure 3B). GapA has been identified as a target of Ag and Cu poisoning in *E. coli* (38) and *Staphylococcus aureus* (39), respectively, and as such, it is a likely candidate for Cu poisoning in GAS. As expected, Cu treatment (both 1 and 5 µM of added Cu) led to a decrease in GapA activity in Δ*copA* mutant cells (Figures 3C(ii), 3D(ii)), which would explain the corresponding reduction in lactate secretion (Figure 3A) and ATP production (Figure 2C). The reduction in GapA activity would also cause upstream glycolytic precursors to accumulate, with consequent feedback inhibition of downstream enzymes (40), as well as glucose phosphorylation and uptake (41, 42) (Supporting Figures 3B(ii), 3D(i)).

This Cu-dependent inhibition is specific to GapA since there was no reduction in GapN activity (Figures 3C(iii), 3D(iii)). Given that there was no detectable change in GapA protein levels in cell extracts (Supporting Figure 4A), these observations are consistent with mismetalation of GapA, as established recently for the GapA homologue in *S. aureus* (39). The excess Cu ions likely bind to the conserved Cys and His residues at the catalytic site (Supporting Figure 4B), as suggested previously for the binding of Ag ions to GapA from *E. coli* (38).

Remarkably, when cultures were sampled earlier (at t = 2 and 3 h), no difference was observed between GapA activity in Cu-treated and control Δ*copA* cells (Figures 3E). The timing of GapA inhibition, i.e. at the onset of the late exponential phase of growth (at t = 4 h; Figure 3E), coincided with the arrest in bacterial growth and metabolism, supporting our hypothesis that GapA is a key target of Cu intoxication in GAS.

### Cu treatment leads to misregulation of metal homeostasis in late exponential phase of growth

The puzzling but consistent, 4-hour delay in the onset of all observable phenotypes led us to hypothesise that there was a time-dependent shift in Cu handling by GAS. To test this proposal, we measured the response of the Cu sensor CopY by monitoring expression of *copZ* during growth in the presence of the lowest inhibitory concentration of added Cu (0.5 µM; see Supporting Figure 2A(ii)). The results show that *copZ* transcription was upregulated ∼4-fold immediately upon Cu exposure (t = 0 h, in which ∼12 min passed between addition of Cu into the culture, centrifugation, and addition of lysis buffer; Figure 4A). This level of upregulation remained largely unchanged during growth (measured up to 5 h; Figure 4A), even though intracellular Cu levels continued to rise (Supporting Figure 5). Our interpretation of these results is that the CopY sensor was fully metalated and expression of *copZ* was already at (or near) its maximum at t = 0 h post-challenge with added Cu. These data also establish that the *copYAZ* operon is transcriptionally induced before the onset of observable growth defects (hereafter referred to as Cu “stress”).

**Figure 4.**
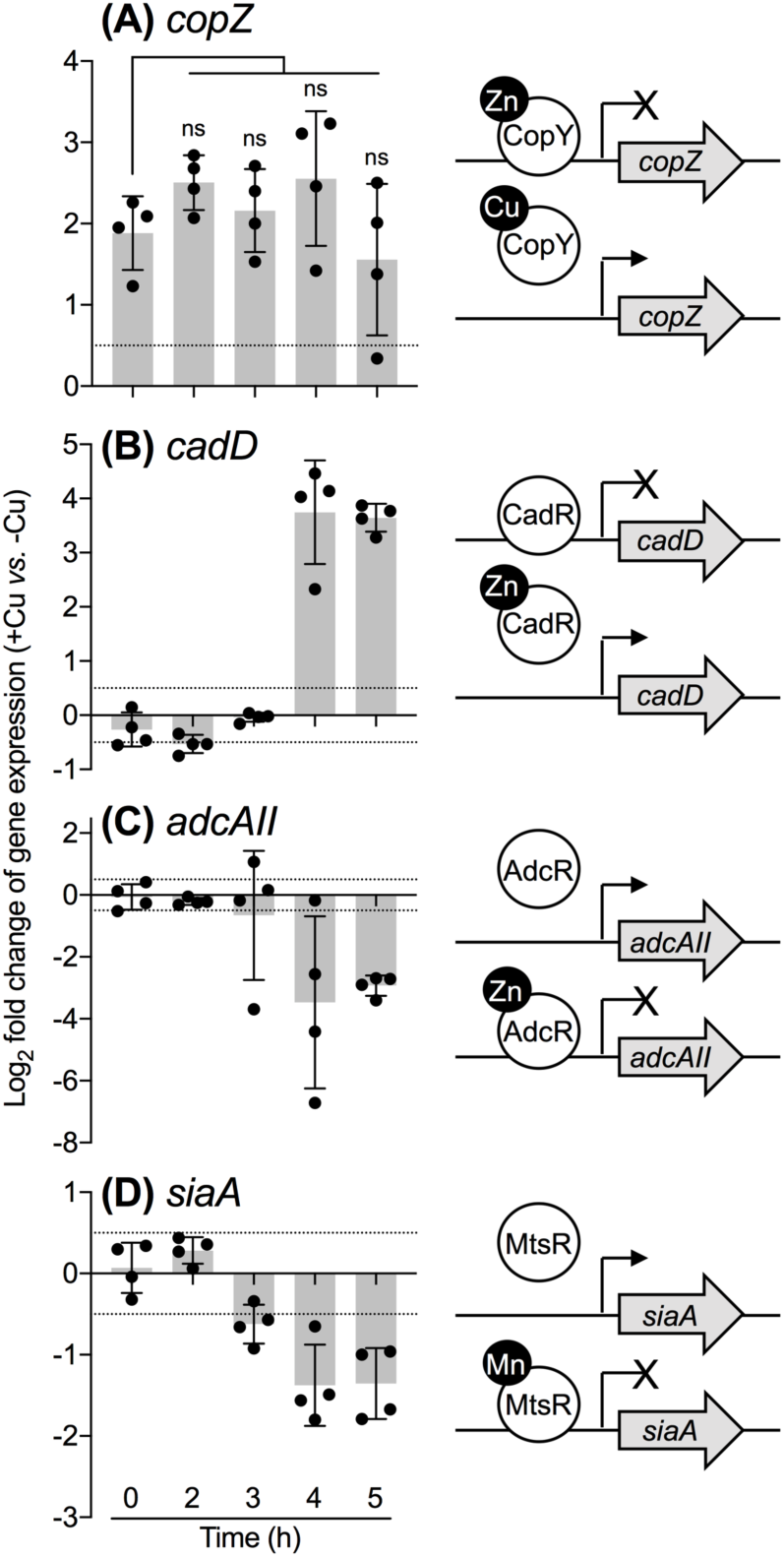
Cu-dependent misregulation of metal homeostasis genes. GAS Δ*copA* mutant strain was cultured with or without added 0.5 µM Cu for the indicated times (*n* = 4). Transcript levels in Cu-treated cultures were determined by qPCR and normalised to the corresponding untreated samples that were cultured for the same time periods. Dotted horizontal lines represent the limit of the assay (log_2_FC = ± 0.5). A schematic representation of each gene and its cognate metallosensor is shown. Transcription of *copZ* or *cadD* is derepressed upon binding of Cu to CopY or Zn (or Cd) to CadR, respectively. Transcription of *adcAII* or *siaA* is repressed upon binding of Zn to AdcR or Mn (or Fe) to MtsR, respectively. **(A) *copZ***. Cu treatment clearly induced *copZ* expression t = 0 h (*P* = 0.0037 *vs.* log_2_FC = 0). This magnitude of induction remained unchanged over the growth period (^ns^*P* = 0.53, 0.94, 0.47, 0.90 for t = 2, 3, 4, 5 h, respectively, *vs.* t = 0 h). **(B) *cadD***. Cu treatment upregulated *cadD* expression at t = 4 and 5 h (*P* = 0.0044 and < 0.0001, respectively, *vs.* log_2_FC = 0). **(C) *adcAII***. Cu treatment downregulated *adcAII* expression at t = 4 and 5 h (*P* = 0.035 and 0.0004, respectively, *vs.* log_2_FC = 0). **(D) *siaA***. Cu treatment downregulated *siaA* expression at t = 3, 4, and 5 h (*P* = 0.014, 0.012, and 0.084, respectively, *vs.* log_2_FC = 0).

We concurrently measured the expression of genes that are controlled by other metalloregulators, namely *adcAII* (regulated by AdcR, a MarR-family Zn-sensing transcriptional co-repressor (43)), *siaA* (controlled by MtsR, a DtxR-family Mn/Fe-sensing co-repressor (44)), and *cadD* (regulated by CadC, an ArsR-family Zn/Cd-sensing derepressor (45)). Clear changes in the expression levels of all three genes were detected in response to Cu treatment. While *adcAII* and *siaA* were downregulated, *cadD* was upregulated (Figures 4B-D). Each of these transcriptional responses is consistent with metalation of the corresponding metallosensor (Figures 4B-D), but whether by the cognate metal or by Cu cannot be distinguished *in vivo*. Nevertheless, the hypothesised effect of Cu on these metallosensors was corroborated by results from genome-wide RNA-seq analyses. Multiple AdcR- and MtsR-controlled genes were negatively regulated, while both the CadC-controlled genes were positively regulated in response to 5 µM of added Cu (Table 1, Supporting Table 1). Interestingly, no clear effect on *gczA* or *czcD* expression was detected, suggesting that the metalation status of GczA, a TetR-family Zn-sensing derepressor (46), is not altered by Cu treatment.

**Table 1.** Cu treatment leads to a misregulation of metal homeostasis. The GAS Δ*copA* mutant strain was cultured with or without 5 µM of added Cu for t = 5 h (*n* = 3). Total RNA was extracted, rRNA was depleted, cDNA was generated, and finally sequenced by Illumina. Differential gene expression was determined using DeSeq2 and presented as the fold-change (FC) of expression in the Cu-treated cultures relative to that in the untreated control. Only genes of interest are listed. These are genes regulated by metal-sensing transcriptional regulators CopY, CadC^45^, AdcR^43^, MtsR^44^, and GczA^46^, as well as those that encode components of the putative GSH uptake system^50^. The complete list of differentially regulated genes is provided in Supporting Table 1.

| Gene | MGAS5005 gene product annotation | $\log_2FC$<br>(+Cu/-Cu) | $P_{adj}$ |
| --- | --- | --- | --- |
| **CopY-regulated** |  |  |  |
| *copZ* | copper chaperone | 4.41 | 0.0000 |
| *copA* | copper-exporting ATPase | 4.39 | 0.0000 |
| *copY* | copAB ATPase metal-fist type repressor | 4.49 | 0.0000 |
| **CadC-regulated** |  |  |  |
| *cadD* | cadmium resistance protein | 3.15 | 0.0000 |
| *cadC* | cadmium efflux system accessory protein | 4.00 | 0.0000 |
| **AdcR-regulated** |  |  |  |
| *phtY* | internalin protein | -1.80 | 0.0000 |
| *rpsN* | 30S ribosomal protein S14 | -1.72 | 0.0000 |
| *phtD* | histidine triad protein | -2.51 | 0.0000 |
| *adcAII* | laminin binding protein | -2.37 | 0.0000 |
| *adhE* | acetaldehyde-CoA/alcohol dehydrogenase | -3.32 | 0.0000 |
| **MtsR-regulated** |  |  |  |
| *fhuG* | ferrichrome transporter permease | -1.99 | 0.0000 |
| *fhuB* | ferrichrome transporter permease | -1.81 | 0.0000 |
| *fhuD* | ferrichrome-binding protein | -1.91 | 0.0000 |
| *fhuA* | ferrichrome ABC transporter ATP-binding protein | -1.75 | 0.0000 |
| *nrdI.2* | ribonucleotide reductase stimulatory protein | -1.57 | 0.0006 |
| *nrdE.2* | ribonucleotide-diphosphate reductase subunit alpha | -1.64 | 0.0001 |
| *hupY* | cell surface protein | -2.29 | 0.0000 |
| *hupZ* | hypothetical protein M5005_Spy_0652 | -2.25 | 0.0000 |
| *siaD* | ABC transporter ATP-binding protein | -2.34 | 0.0000 |
| *siaC* | ferrichrome ABC transporter ATP-binding protein | -2.71 | 0.0000 |
| *siaB* | ferrichrome transporter permease | -2.63 | 0.0000 |
| *siaA* | ferrichrome-binding protein | -2.28 | 0.0000 |
| *shp* | heme binding protein | -1.80 | 0.0000 |
| *shr* | Fe<sup>3+</sup>-siderophore transporter | -1.94 | 0.0000 |
| **GczA-regulated** |  |  |  |
| *gczA* | TetR family transcriptional regulator | 0.03 | 0.9378 |
| *czcD* | cobalt-zinc-cadmium resistance protein | 0.46 | 0.2807 |
| **GSH import** |  |  |  |
| *gshT* | amino acid ABC transporter substrate-binding protein | 0.90 | 0.1098 |
| *tcyB* | amino acid ABC transporter permease | 0.83 | 0.0409 |
| *tcyC* | ABC transporter substrate-binding protein | 0.87 | 0.0443 |

Crucially, changes in the expression of *adcAII, siaA*, and *cadD* appeared only after ∼4 h of growth (Figures 4B-D). These transcriptional changes were not accompanied by increases in total intracellular Zn, Mn, or Fe levels (Supporting Figure 5). Thus, the simplest model that accounts for the sudden metalation (or mismetalation) of multiple metallosensors, as well as GapA, is that excess Cu is released from an intracellular buffer, leading to mislocation of Cu to adventitious binding sites and/or redistribution of intracellular metals.

### The onset of the Cu stress phenotype coincides with depletion of GSH

What comprises the intracellular buffer for excess Cu in GAS? This organism does not possess a homologue of the metallothionein MymT (17) or the Cu storage protein Csp (47). Instead, this buffer likely consists of a polydisperse mixture of cytoplasmic small molecules or metabolites (48). Noting that GAS is auxotrophic for most nutrients, including multiple amino acids, vitamins, nucleobases, and GSH, we hypothesised that: (i) one or more of these nutrients constitute the intracellular Cu buffer, either directly by coordinating Cu or indirectly by acting as a synthetic precursor to the buffer; and that (ii) these nutrients become exhausted from the extracellular medium during bacterial growth, leading to the observable effects of Cu stress.

We tested this hypothesis using two complementary approaches and the results identify GSH as the key limiting nutrient. First, mass spectrometry was employed to measure consumption of nutrients from the growth medium. Several amino acids, the nucleobases adenine and uracil, as well as GSH (and/or its disulfide GSSG) were nearly or completely spent after ∼4 h of growth (Supporting Figure 6). Cys and its disulfide were below detection limits. Next, the culture medium was supplemented with each or a combination of the spent or undetected extracellular nutrients. Their ability to restore growth of the Δ*copA* mutant in the presence of added Cu was subsequently examined. Only supplementation with GSH was strongly protective against Cu intoxication (Supporting Figure 7).

The GAS genome encodes neither the common pathway for GSH biosynthesis (GshAB) nor the bifunctional glutathione synthetase (GshF (49)). Instead, an uncharacterised homologue of the GSH-binding solute-binding protein GshT is present (Supporting Figure 8). GshT, in conjunction with the endogenous cystine importer TcyBC, likely allows GAS to import extracellular GSH (γ-Glu-Cys-Gly) into the cytoplasm (50). This system may also transport γ-Glu-Cys or Cys-Gly (50), but addition of these dipeptides, or Cys alone, or a mixture of the amino acids Glu, Cys, and Gly did not improve growth of Cu-treated Δ*copA* mutant cultures (Figure 5A). Altogether, these results suggest that: (i) the protective effect of GSH is unlikely to result from chelation of *extracellular* Cu ions by thiophilic ligands; (ii) extracellular GSH is depleted during growth of GAS; and (iii) this depletion is responsible for the observable Cu stress phenotypes. Consistent with propositions (ii) and (iii), addition of GSH completely suppressed the effects of Cu treatment and restored plating efficiency, as well as glucose consumption, lactate secretion, and ATP production beyond the late exponential phase of growth (Figures 5B-E).

**Figure 5.**
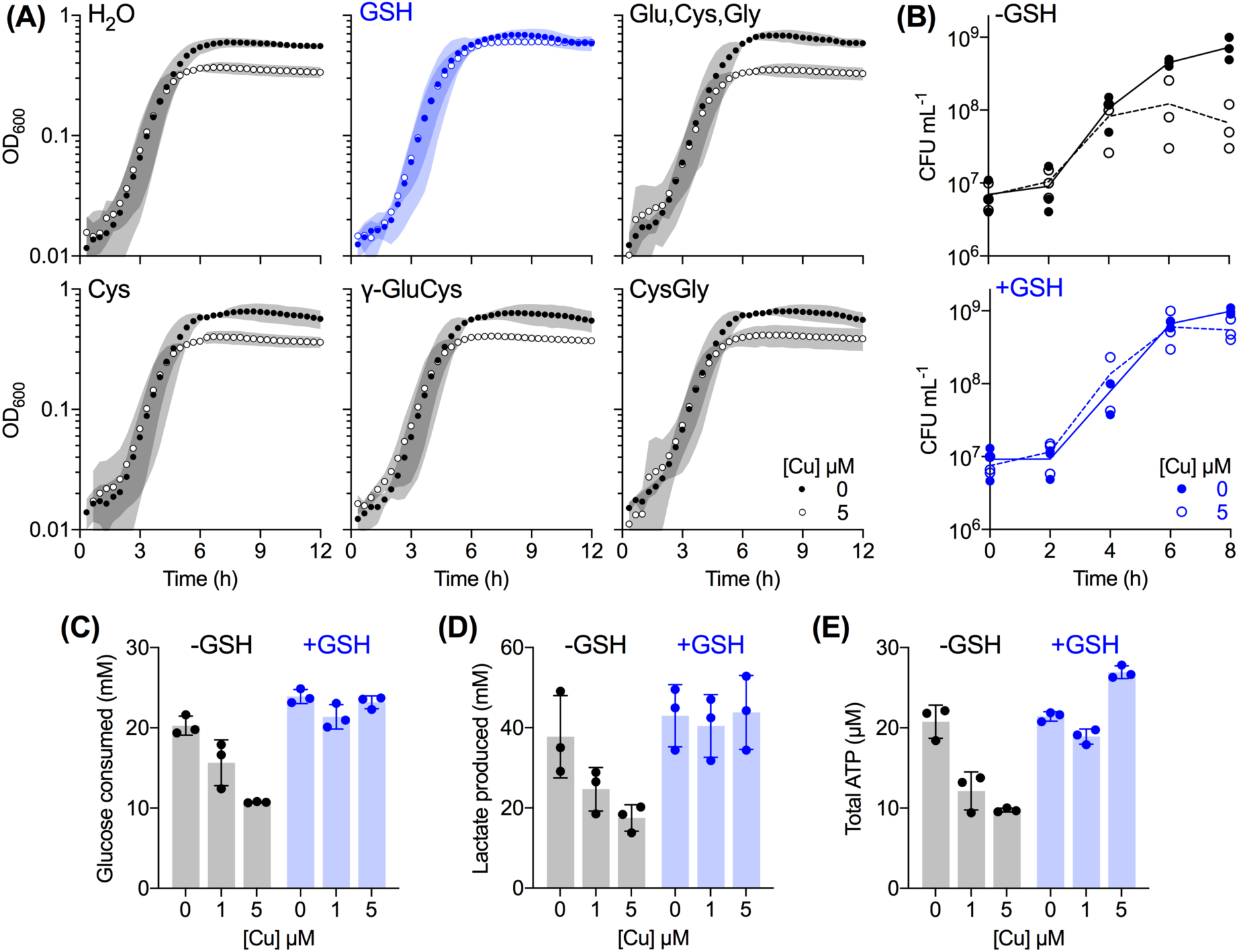
Protective effects of supplemental GSH. **(A) Growth.** GAS Δ*copA* mutant strain was cultured with added Cu as indicated (*n* = 3). The culture medium was also supplemented with: water or 0.1 mM each of GSH (blue); a mixture of Glu, Cys, and Gly; Cys alone; the dipeptide γ-GluCys; or CysGly. Cu treatment did not affect GSH-supplemented cultures (*P* = 0.99). **(B)-(E)** GAS Δ*copA* mutant strain was grown with added Cu as indicated, without (black) or with (blue) 0.1 mM GSH (*n* = 3). **(B) Plating efficiency.** Cultures were plated out at the indicated time points and the numbers of colony-forming units (CFU) were enumerated. Cu treatment clearly suppressed plating efficiency of GSH-deplete cultures (*P* = 0.0012) but not that of the GSH-supplemented cultures (*P* = 0.97). **(C) Glucose consumption.** Cultures were sampled at t = 8 h and total amounts of glucose consumed from the extracellular growth media were determined. Cu treatment clearly suppressed glucose consumption by GSH-deplete cultures (*P* = 0.0053 for 1 µM Cu, *P* < 0.0001 for 5 µM Cu) but not that by GSH-supplemented cultures (*P* = 0.12 for 1 µM Cu, *P* = 0.81 for 5 µM Cu). **(D) Lactate production**. Cultures were sampled at t = 8 h and amounts of lactate secreted to the extracellular growth media were determined. Cu treatment clearly suppressed lactate production by GSH-deplete cultures (*P* = 0.11 for 1 µM Cu, *P =* 0.014 for 5 µM Cu) but not that by GSH-supplemented cultures (*P* = 0.91 for 1 µM Cu, *P =* 0.99 for 5 µM Cu). **(C) Total ATP levels.** Cultures were sampled at t = 8 h and total ATP levels were determined. Cu treatment clearly suppressed ATP production by GSH-deplete cultures (*P* < 0.0001 each for 1 and 5 µM Cu) but not that by the GSH-supplemented cultures (*P* = 0.095 for 1 µM Cu, *P =* 0.0008 for 5 µM Cu). All statistical analyses were *vs.* 0 µM Cu.

### GSH contributes to buffering of excess intracellular Cu

The time-dependent reduction in extracellular GSH levels (Supporting Figure 6D) was mirrored by a decrease in intracellular GSH (Figure 6A). Both the wild type and Δ*copA* mutant strains contained ∼4 mM of intracellular GSH (and GSSG) at t = 0 h (Figure 6A). This amount was likely already present in the inoculum, which was cultivated in the complex medium THY ([GSH]_THY_ ∼ 30 µM (51)). Intracellular GSH levels in both strains reduced to ∼0.1 mM at t = 4 h, regardless of Cu treatment (Figures 6A-B). This decrease occurred presumably as a consequence of bacterial growth and replication in a chemically defined medium with a limited extracellular supply of this thiol ([GSH]_CDM_ ∼ 0.5 µM; Supporting Figure 6D). This low amount of GSH coincided with the onset of the observable Cu stress phenotypes. It might also explain why cultures that grew to low OD_600_ values displayed no sign of Cu stress (Supporting Figures 2C-D) – these cultures likely had not depleted their intracellular GSH supply.

**Figure 6.**
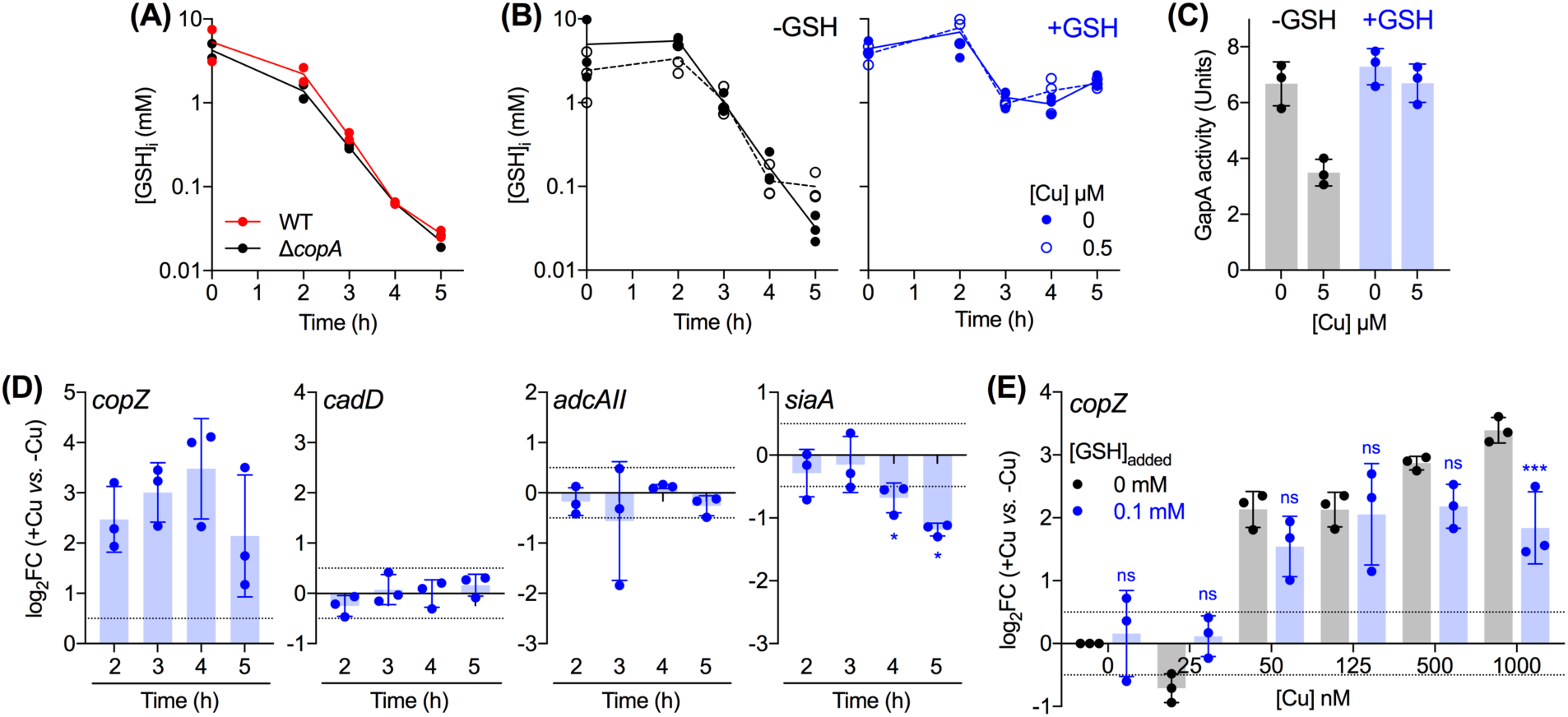
Buffering of excess intracellular Cu ions by GSH. **(A) Time-dependent changes in intracellular GSH concentrations.** GAS strains were cultured without any added Cu or GSH (*n* = 2). Cultures were sampled at the indicated time points and intracellular levels of GSH were measured in cell extracts. There was no clear difference between the intracellular GSH levels of WT and Δ*copA* cultures (*P* = 0.09). **(B)-(E) Effects of GSH supplementation.** GAS Δ*copA* mutant strain was cultured with added Cu as indicated, without (black) or with 0.1 mM of added GSH (blue). **(B) Intracellular GSH concentrations.** Cultures (*n* = 3) were sampled at the indicated time points. Intracellular levels of GSH were measured in cell extracts. Cu treatment did not affect intracellular GSH levels, regardless of GSH supplementation (*P* = 0.95 for 0 mM GSH, *P* = 1.0 for 0.1 mM GSH). GSH supplementation clearly improved intracellular GSH levels (*P* < 0.0001 for both 0 and 0.5 µM Cu), regardless of Cu treatment. **(C) GapA activity.** Cultures (*n* = 3) were harvested at t = 4 h. GapA activity was measured in cell extracts. Cu treatment had a clear effect on GapA activity in GSH-deplete cultures (*P* = 0.0007) but not on GSH-supplemented cultures (*P* = 0.51). **(D) Expression of metal homeostasis genes.** GSH-supplemented cultures (*n* = 3) were sampled at the indicated time points. Levels of *copZ, cadD, adcAII*, and *siaA* transcripts in Cu-treated cultures were determined by qPCR and normalised to the corresponding untreated samples that were harvested at the same time points. Horizontal dotted lines represent the limit of the assay (log_2_FC = ± 0.5). Cu treatment induced *copZ* expression (*P* = 0.023, 0.013, 0.026, 0.093), but not *cadD* (*P =* 0.17, 0.71, 0.98, 0.32), or *adcAII* (*P* = 0.39, 0.50, 0.03, 0.16) at t = 2, 3, 4, 5 h, respectively (*vs.* log_2_FC = 0). Cu treatment continued to downregulate *siaA* expression (*P* = 0.63, 0.03, 0.04, 0.03 for t = 2, 3, 4, 5 h, respectively, *vs.* log_2_FC = 0). **(E) Cu-dependent expression of *copZ***. Cultures (*n* = 3) were sampled at t = 4 h. Levels of *copZ* transcripts in Cu-treated cultures were normalised to the corresponding untreated samples. Horizontal dotted lines represent the limit of the assay (log_2_FC = ± 0.5). GSH supplementation did not affect *copZ* expression at low concentrations of added Cu (^ns^*P* = 0.10, 0.14, 0.48, 1.0, and 0.31 for [Cu] = 0, 25, 50, 125, and 500 nM) but it did affect expression at 1000 nM of added Cu (\*\*\**P* = 0.0009).

We noted that Cu treatment did not transcriptionally induce the uptake of GSH. Levels of *gshT* transcripts remained largely unchanged, based on RNA-seq analyses of *ΔcopA* cells at the late-exponential phase of growth (Table 1). This result supports previous transcriptomic studies in several Gram-positive and Gram-negative bacteria, none of which identified GSH biosynthesis or uptake as a key transcriptional response to Cu treatment (15, 20, 22, 23).

Supplementation of the growth medium with GSH (0.1 mM) did not affect the intracellular levels of this thiol at the early stages of growth (t = 0 and 2 h; Figure 6B). However, it did allow Δ*copA* cells to maintain intracellular concentrations of this tripeptide at ∼1 mM (one log unit higher than unsupplemented cells) beyond the late exponential growth phase, regardless of Cu treatment (Figure 6B). As mentioned earlier, these GSH-treated cells were Cu-tolerant (Figure 5).

A more detailed examination of GSH-supplemented Δ*copA* cells revealed that GapA was protected from inactivation by added Cu (Figure 6C). In addition, the Cu-induced, time-dependent changes in *cadD* and *adcAII* expression were abolished (Figure 6D), suggesting that CadC and AdcR were protected from mismetalation. We continued to observe some downregulation of *siaA* transcription, albeit to a lesser magnitude when compared with GSH-deplete cultures (Figure 6D *vs.* Figure 4D). In general, these results support a model whereby GSH constitutes the major buffer for excess intracellular Cu in GAS and protects potential non-cognate binding sites from becoming (mis)metalated by Cu.

Importantly, GSH supplementation did not affect expression of *copZ* at low concentrations of added Cu (0 – 500 nM; Figure 6E). This observation is consistent with the proposal that GSH does not rescue the Δ*copA* mutant simply by chelating extracellular Cu ions. However, GSH treatment did partially suppress *copZ* expression in response to a high concentration of added Cu (1000 nM; Figure 6E). This observation indicates the relative buffering strengths of GSH and CopY, which are discussed below.

## DISCUSSION

### The role of GSH in buffering of excess cytoplasmic Cu

GSH has been proposed to bind Cu by assembling a stable, tetranuclear Cu_4_GS_6_ cluster (52). In such a model, when present at low millimolar concentrations (e.g. ∼4 mM in GAS at t = 0 h, see Figure 6A), GSH would bind Cu with an apparent affinity of *K*D = 10^−16.7^ M and thus would impose a threshold of Cu availability at 10^−16.7^ M (Supporting Figure 9A). This threshold is above the range of Cu availability set by most bacterial Cu sensors (Supporting Figure 9B) (53–55). Therefore, GSH contributes to Cu buffering only when the transcriptionally-responsive Cu homeostasis system is impaired (e.g. in a Δ*copA* mutant (25, 26)) or overwhelmed (e.g. when intracellular Cu levels rise above the responsive range of the Cu sensors).

Figure 6E shows that supplementation with GSH had little impact on metalation of CopY (and thus expression of *copZ*) when the amounts of added Cu were low. However, GSH appeared to dampen the CopY response at higher concentrations of added Cu, indicating that this thiol competes with CopY for binding Cu when intracellular Cu levels are high. Hence, the thresholds of intracellular Cu availability set by GSH and CopY may overlap, at least partially, with GSH being the weaker buffer (52, 55, 56). The model in Supporting Figure 9B is compatible with these experimental data but it will need refinement. This model was estimated using known parameters (Cu affinity, DNA affinity, number of DNA targets) for CopY from *S. pneumoniae* (CopY_Spn_) (55), but CopY_Spn_ differs from CopY_GAS_ in several key aspects. CopY_Spn_ lacks one of the two Cys-X-Cys motifs found in other CopY homologues such as CopY_GAS_ and CopY from *E. hirae* (CopY_Eh_) (Supporting Figure 10). CopY_Spn_ binds 2 Cu atoms per dimer in a solvent-exposed centre while CopY_Eh_ binds 4 Cu atoms per functional dimer and assembles a solvent-occluded centre (55, 57). In addition, two *cop* boxes are present in *S. pneumoniae* (58) while only one is found in GAS. How these differences shift the threshold model will need to be examined using careful *in vitro* studies with purified proteins and DNA. In the simplest scenario, an increase in the stability (affinity) of the bound Cu atoms in CopY, which may occur as a consequence of coordination by extra Cys ligands, would lower the threshold of Cu availability set by CopY (Supporting Figure 9C), and thus better fit our experimental data.

Depletion of intracellular GSH to 0.1 mM at the late exponential phase of growth would weaken its buffering capacity by at least 2 log units (Supporting Figure 9A). Figure 4 shows that Cu is then able to metalate non-specific binding sites in non-cognate metallosensors or metalloenzymes. Our results further suggest that AdcR, CadC, and MtsR can allosterically respond to Cu and differentially regulate expression of their target genes *in vivo*. Precisely how this occurs will need to be confirmed with purified proteins *in vitro.* Cu-responsive regulation of genes under control of non-cognate metallosensors has indeed been reported both *in vivo* and *in vitro*, although not for the families of regulators described here (15, 28, 59–61).

Not all bacteria use GSH as the major cytoplasmic thiol. Some bacilli, such as *B. subtilis* and *S. aureus*, produce the glycoside bacillithiol (BSH) instead. The affinity of BSH to Cu is at least 2 orders of magnitudes tighter than that of GSH (56, 62). Hence, BSH likely imposes a lower limit on cytoplasmic Cu availability than does GSH but it is worth noting that its intracellular level is ∼30 times lower than that of GSH (63). Importantly, the relative order with the endogenous Cu sensor CsoR still holds, with BSH binding Cu at least 3 log units more weakly than does CsoR (54). Indeed, this thiol is also thought to contribute to Cu homeostasis by buffering excess Cu. Deletion of the *B. subtilis bshC* gene for BSH biosynthesis led to a slight increase in *copZ* expression in response to added Cu. This result mirrors our finding in Figure 6E and suggests that the Cu sensor CsoR is more readily metalated by Cu in the absence of the major buffering thiol (64). It is also notable that the identification of GapA as a major reservoir of excess Cu ions in the cytoplasm was in a strain of *S. aureus* that does not synthesise BSH (39).

In summary, this study provides a new line of evidence that Cu handling in the bacterial cytoplasm, when formulated using the threshold model, comprises two components (Figure 7). The transcriptionally-responsive component, which includes the Cu sensor, Cu efflux pump, and additional Cu-binding metallochaperones, functions in housekeeping or *homeostasis*, and sets a low limit of Cu availability in the cytoplasm. Rising Cu levels can saturate this homeostasis system and sudden Cu shock can overwhelm it, but the transcriptionally-unresponsive component, in this case GSH, buffers the excess Cu and confers additional Cu *tolerance*. This second system acts as the final layer of protection before cells experience widespread mismetalation and, therefore, Cu *stress* (Figure 7). This additional buffering essentially extends the range of cytoplasmic Cu availability that can be tolerated by bacteria, allows bacteria to maintain key cellular functions, and thus prevents an abrupt transition from Cu homeostasis to Cu stress upon exposure to an excess of this metal ion.

**Figure 7.**
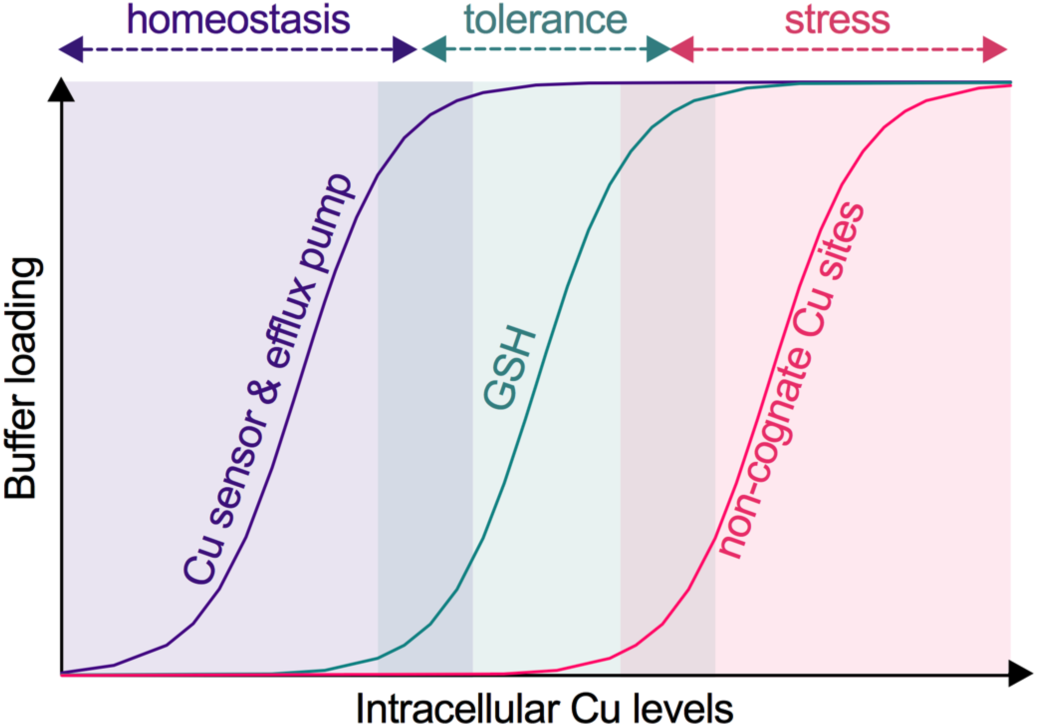
Threshold model for bacterial Cu homeostasis, tolerance, and stress. As Cu levels in the cytoplasm increase, this metal ion binds to the allosteric site of the Cu-sensing transcriptional regulator, which subsequently induces expression of the Cu efflux pump. Together, the Cu sensor and efflux pump impose a low limit of Cu availability and maintain Cu homeostasis. A further rise in cytoplasmic Cu levels saturates the Cu homeostasis system and begins to fill binding sites in GSH. Since there are no observable defects in bacterial metabolism or growth at this stage, GSH can be considered to confer Cu tolerance. GSH depletion or a further increase in cytoplasmic Cu levels saturates this tolerance capacity. Cu now binds to non-cognate metal-binding sites, leading to inhibition of bacterial metabolism and growth. These conditions are considered as Cu stress.

### The role of GSH in buffering bacterial Cu during host-pathogen interactions

This study was conducted originally to examine the role of the Cop Cu homeostasis system in GAS pathogenesis. Although GAS occupied a Cu-rich environment in mice (Figure 1A), inactivation of the *copA* gene did not lead to a reduction in GAS virulence (Figure 1C). Our *in vitro* investigations now suggest that GAS may withstand host-imposed increases in Cu levels, as long as it has access to a source of GSH *in vivo*. Indeed, we did detect the presence of GSH in the skin ulcers of infected mice, but interestingly, the amount was ∼25-fold less when compared with skin from healthy mice or healthy skin from infected mice (Supporting Figure 11). Whether this depletion of GSH is a feature of the general host immune response, a consequence of inflammation and/or host tissue necrosis, or a consequence of GAS metabolism is not known. Nevertheless, the virulence of the Δ*copA* mutant implies that the level of host GSH, albeit reduced, can support Cu buffering inside the GAS cytoplasm. Alternatively, the level of host Cu (Figure 1A) may not be sufficient to overwhelm the Cop homeostasis system, since the *copYAZ* operon was only slightly upregulated in bacteria isolated from mouse ulcers (average log_2_FC = 1.13 *vs.* THY) (34).

### The link between the failure to buffer Cu and redox stress

Under our experimental conditions, untreated Δ*copA* cells contained 20,000 (± 3,000) Cu atoms when sampled at t = 3 h (before the onset of Cu stress). Cu treatment increased this number ∼10-fold to 190,000 (±130,000) atoms (Supporting Figure 5). The intracellular GSH concentrations at the same time point ([GSH]_i_ = 0.76 mM, Figure 6A) would translate to ∼500,000 molecules of GSH, which are clearly insufficient to buffer all of the intracellular Cu ions. Yet, there was no observable Cu stress phenotype at this time point, suggesting that the excess Cu remains bound to other cytoplasmic component(s). These components may include CopZ and/or the novel, uncharacterised protein CopX (Supporting Figure 1A). This idea will be the focus of future studies.

Finally, the GSH/GSSG couple is the major redox buffer of the cell. Assuming that the GSH/GSSG ratio remains unchanged, depletion of intracellular GSH in GAS from ∼4 mM to ∼0.1 mM would raise the cytoplasmic redox mid-potential by ∼46 mV. This relatively more oxidising environment, when combined with a lack of Cu buffering, may promote Cu-catalysed generation of reactive oxygen species (65) or formation of disulfides (66). Yet, our RNA-seq results do not suggest widespread oxidative stress (Supporting Table 1). In *E. coli*, deletion of *gshA* did not accelerate DNA damage in Cu-replete cells, even in the presence of added H_2_O_2_ (67). Similarly, proteomic analyses of a non-BSH producing strain of *S. aureus*, indicated that Cu-treatment does not induce a strong oxidative stress response in this organism (39).

Regardless of the relative importance of mismetalation *vs.* redox stress, our work demonstrates that excess Cu is not bacteriotoxic as long as cytoplasmic GSH is abundant and thus able to buffer the excess of this metal ion (Figure 7). In GAS, a GSH auxotroph, this intracellular buffer is dynamic; its levels change during bacterial growth and/or in response to extracellular GSH availability. Future studies should take these effects into account when examining the impact of Cu treatment on bacterial cultures. Had our work not identified the 4 h time point as metabolically relevant, sampling cultures 1 h earlier would have led to a different conclusion.

## METHODS

### Data presentation and statistical analyses

We follow recent recommendations regarding transparency in data representation (68, 69). Except for growth curves, individual data points from independent experiments are plotted, with shaded columns representing the means, and error bars representing standard deviations. Growth curves show the means of independent experiments, with shaded regions representing standard deviations. The number of independent experiments is stated clearly in each figure legend. Statistical analyses have been performed on all numerical data but *P* values are displayed on plots only if they aid in rapid, visual interpretation. Otherwise, *P* values for key comparisons are stated in the figure legends. Unless otherwise stated, statistical tests used two-way ANOVA using the statistical package in GraphPad Prism 8.0. All analyses were corrected for multiple comparisons.

### Ethics statement

Animal experiments were conducted according to the Guidelines for the Care and Use of Laboratory Animals (National Health and Medical Research Council, Australia) and were approved by the University of Queensland Animal Ethics Committee (Australia). Human blood donation for use in neutrophil killing studies was conducted in accordance with the National Statement on Ethical Conduct in Human Research and in compliance with the regulations governing experimentation on humans, and was approved by the University of Queensland Medical Research Ethics Committee (Australia).

### Reagents

All reagents were of analytical grade and obtained from Sigma or Melford Chemicals unless otherwise indicated. γ-Glu-Cys and Cys-Gly were from BACHEM Peptides (Germany). The sulfate and chloride salts of copper were used interchangeably. All reagents were prepared in deionised water.

### Strains and culture conditions

GAS M1T1 5448 strains were propagated from frozen glycerol stocks onto solid THY medium without any antibiotics. Unless otherwise indicated, liquid cultures were prepared in a chemically defined medium containing glucose as the carbon source (CDM-Glucose; Supporting Table 2). This medium routinely contained 53 nM basal Cu, 155 nM Zn, 66 nM Fe, 9 nM Mn, 29 nM Co, and 23 nM Ni, as determined by ICP MS. All solid and liquid growth media contained catalase (50 µg/mL).

### Construction of mutants

Non-polar GAS mutant strains were constructed by allelic exchange following standard protocols (70). Primers and plasmids used in this study are listed in Supporting Tables 3 and 4, respectively. All constructs and genetically altered strains were confirmed by PCR and Sanger sequencing.

### Mice virulence assays

Transgenic, human plasminogenised *AlbPLG1* mice heterozygous for the human transgene were backcrossed greater than *n* = 6 with C57BL/J6 mice as described previously (71). GAS was prepared to obtain the target dose in the 10^7^ CFU range (WT 1.8×10^7^, Δ*copA* 1.5×10^7^) immediately prior to injection. Mice were subcutaneously infected (*n* = 10) and virulence was determined by observing survival for 10 days post-infection. Metal levels in mouse blood and skin were measured by ICP MS as described previously (72).

To assess GSH levels at the site of infection, mouse skin and infected lesions were excised 3 days post-infection, washed with PBS, resuspended in 1 mL PBS, homogenised in Lysing Matrix F tubes using a FastPrep 24G instrument (MP Biomedicals, 4^°^C, speed 6, 40 s, 2 cycles), and centrifuged (10,000 x *g*, 5 min, 4°C). Total GSH was measured from the supernatant using the GSH-Glo kit (Promega) following manufacturer’s instructions, with the modification of mixing undiluted samples 1:1 with 2 mM of tris(2-carboxyethyl)phosphine (TCEP) immediately prior to use.

### Neutrophil killing assays

Survival of GAS following incubation with human neutrophils *ex vivo* was assayed at a multiplicity of infection of 10:1 as previously described (71).

### Bacterial growth

Growth was assessed at 37°C in flat-bottomed 96-well plates using an automated microplate shaker and reader. Each well contained 200 µL of culture. Each plate was sealed with a gas permeable, optically clear membrane (Diversified Biotech). OD_600_ values were measured every 20 min for 12 h. The plates were shaken at 200 rpm for 1 min in the double orbital mode immediately before each reading. OD_600_ values were not corrected for path length (ca. 0.58 cm for a 200 µL culture).

### Plating efficiency

GAS was cultured in 96-well plates as described earlier for growth analysis, sampled at the indicated time points, vortexed for 30 s, diluted serially in PBS, and plated onto solid THY medium without any antibiotics. Colony forming units (CFUs) were enumerated after overnight incubation at 37°C.

### ATP levels

GAS was cultured in 96-well plates as described earlier for growth analysis and sampled at the indicated time points. The amount of total ATP in each sample was determined immediately using the BacTiter-Glo kit (Promega).

### Intracellular metal content

GAS was cultured in 10 – 500 mL of CDM-Glucose as required (larger volumes were required to obtain enough biomass at earlier time points). At the desired time points, an aliquot was collected for the measurement of OD_600_ or plating efficiency. The remaining cultures were harvested (5,000 x *g*, 4°C, 10 min), washed once with PBS containing EDTA (1 mM) and twice with ice-cold PBS. The final pellet was dissolved in concentrated nitric acid (150 µL, 80°C, 1 h) and diluted to 10 mL with deionised water. Total metal levels were determined by ICP MS. The results were normalised to OD_600_ values or plating efficiency as indicated in the figure legends.

### Fermentation end products

GAS was cultured in 96-well plates as described earlier for growth analysis. At the desired time points, samples were centrifuged (5,000 x *g*, 4°C, 10 min) and the supernatants were frozen at -20°C until further use. Concentrations of pyruvate, lactate, acetate, and ethanol in the spent culture media were determined using K-PYRUV, K-LATE, K-ACET, and K-ETOH kits (Megazyme), respectively. Concentrations of glucose were determined using the GAGO20 kit (Sigma).

### Enzyme activity

GAS was cultured in 40 – 250 mL of CDM-glucose as required (larger volumes were required to obtain enough biomass at earlier time points). At the desired time points, bacteria were harvested (5,000 x *g*, 4°C, 10 min), washed once with PBS, and frozen at -20°C until further use. Bacterial pellets were resuspended in a buffer containing sodium phosphate (100 mM) and triethanolamine (80 mM) at pH 7.4, transferred to a tube containing Lysing Matrix B (MP Biomedicals), and lysed in a FastPrep 24G instrument (MP Biomedicals, 10 m/s, 20 s, 2 cycles). Cell debris were removed by centrifugation (20,000 x *g*, 1 min). The cell-free lysate supernatant was kept on ice and used immediately.

To determine GapA activity, the reaction mixture contained NAD^+^ (4 mM), *DL-*glyceraldehyde-3-phosphate (G3P, 0.3 mg/mL), sodium phosphate (100 mM), DTT (1 mM), and triethanolamine (80 mM) at pH 7.4. GapN activity was determined as above for GapA but using NADP^+^ (4 mM) instead of NAD^+^ as the electron acceptor. To measure the activity of Ldh, the reaction mixture contained NADH (4 mM), pyruvate (10 mM), and fructose-1,6-bisphosphate (1 mM) in PBS at pH 7.4. For all three enzymes, each reaction (100 µL) was initiated by addition of cell-free extracts (10 µL). Absorbance values at 340 nm were monitored for up to 10 min at 37°C. The initial rates of reaction were normalised to total protein content as determined using the QuantiPro BCA Assay Kit (Sigma). Control reactions without any substrate (G3P for GapA and GapN, pyruvate for Ldh) were always performed in parallel.

One unit of activity was defined as follows: 1000 nmol NAD^+^ oxidised min^-1^ mg protein^-1^ for GapA, 100 nmol NADP^+^ oxidised min^-1^ mg protein^-1^ for GapN, and 1000 nmol NADH reduced min^-1^ mg protein^-1^ for Ldh.

### GSH levels

GAS was cultured in 10 – 150 mL of CDM-glucose as required (larger volumes were required to obtain enough biomass at earlier time points). At the desired time points, an aliquot was plated for bacterial counting. The remaining cultures were harvested (5,000 x *g*, 4°C, 10 min), washed twice with PBS, resuspended in 5-sulfosalycylic acid (5 w/v %), transferred to a tube containing Lysing Matrix B, and frozen at -20°C until further use. Bacteria were lysed in a bead beater (10 m/s, 30 s, 2 cycles). Cell debris were removed by centrifugation (20,000 x *g*, 1 min). Total GSH (and GSSG) levels in lysate supernatants were determined immediately using the Gor-DTNB recycling method (73) and normalised to total bacterial counts.

### RNA extraction

GAS was cultured in 2 – 200 mL of CDM-glucose as required (larger volumes were required to obtain enough biomass at earlier time points). At the desired time points, cultures were centrifuged (3,000 x *g*, 4°C, 5 min). Bacterial pellets were resuspended immediately in 1 mL of RNAPro Solution (MP Biomedicals) and stored at -80°C until further use. Bacteria were lysed in Lysing Matrix B and total RNA was extracted following the manufacturer’s protocol (MP Biomedicals). RNA extracts were treated with RNase-Free DNase I enzyme (New England Biolabs). Complete removal of gDNA was confirmed by PCR using gapA-check-F/R primers (Supporting Table 3). gDNA-free RNA was purified using RNeasy Mini Kit (QIAGEN) and visualised on an agarose gel.

### qPCR analyses

cDNA was generated from 1 µg of RNA using the SuperScript® IV First-Strand Synthesis System (Invitrogen). qPCR was performed in 10 or 20 µL reactions using 2 or 5 ng of cDNA as template and 0.4 µM of the appropriate primer pairs (Supporting Table 3). Each sample was analysed in technical duplicates. Amplicons were detected with PowerUP SYBR Green (Invitrogen) in a QuantStudio 6 Flex Real-Time PCR System (Applied Biosystems) or a CFXConnect Real-Time PCR Instrument (Bio-Rad Laboratories). *C*q values were calculated using LinRegPCR after correcting for amplicon efficiency. *C*q values of technical duplicates were typically within ± 0.25 of each other.

*holB* and *tufA*, which encode DNA polymerase III and elongation factor Tu, respectively, were used as reference genes (Supporting Table 3). Their transcription levels remained constant in all of the experimental conditions tested here. *holB* was used as the reference gene in all the data presented here because its *C*q values were closer to the dynamic ranges of *cop* genes, *adcAII, cadD*, and *siaA*, but the results were identical with when *tufA* was used as the reference.

### RNA-seq analyses

GAS Δ*copA* mutant strain was cultured in the presence of 0 or 5 µM of added Cu for t = 5 h (*n =* 3) and RNA was extracted from each culture as described earlier. RNA-Seq was performed from Ribo-zero (rRNA depleted) triplicate samples on a single Illumina HiSeq 2500 lane using v4 chemistry from 75 base pair paired-end reads. Reads were mapped to the 5448 (M1) GAS reference genome (GenBank accession number CP008776.1) with BWA MEM (version 0.7.16). Relative read counts (per gene) and differential gene expression was determined using DESeq2 (v. 1.26.0) (74) in R. Genes with less than 10 reads across all conditions and samples were removed. *P*-values were calculated using Wald test and adjusted for multiple testing using Benjamini-Hichberg/false discovery rate. Illumina read data were deposited to the European Nucleotide Archive Sequence Read Archive under the accession numbers ERS1996831, ERS1996835, and ERS1996839.

## AUTHOR CONTRIBUTIONS

AGM, KJW, KYD, MJW initiated the research. KYD had overall responsibility for the conceptualisation and coordination of the programme. KYD designed the experiments with input from KJW. C-LYO, MMZ generated the Δ*copA* and *copA*^+^ mutant strains. KYD, LJS conducted the *in vitro* experiments. C-LYO, MMZ performed infection assays in neutrophils. C-LYO, MZ, SB performed mice infection assays. MRD, LM conducted the RNA-seq analyses. All authors contributed to data analysis. KJW, KYD, LJS wrote the initial manuscript. All authors reviewed and approved the final version of the manuscript.

## ACKNOWLEDGEMENT

We thank members of the Metals in Biology grouping (Department of Biosciences, Durham University) for helpful discussions related to this project and Dr Robert Borthwick (Department of Physics, Durham University) for reviewing the manuscript. Dr Marietjie Mostert (School of Earth Sciences, The University of Queensland) and Dr Deenah Morton (Department of Biosciences, Durham University) provided technical assistance with ICP MS. Dr Amanda Walker and Dr Nadia Keller (School of Chemistry and Molecular Biosciences, The University of Queensland) assisted with collection of mice tissues. Dr Ian Cummins (Department of Biosciences, Durham University) provided technical assistance with measurements of growth media components using mass spectromerty. We acknowledge the assistance of the sequencing and pathogen informatics core teams at the Wellcome Trust Sanger Institute.

