## Supporting Tables 3-4 for "A role for glutathione in buffering excess intracellular copper in *Streptococcus pyogenes*"

**Supporting Table 3. List of primers used in this study.** Primers were designed using the genome sequence from *S. pyogenes* MGAS5005 as template (NCBI GenBank Accession No CP000017.2). Primers were purchased from Sigma.

| Amplicon name | Primer name | Sequence | Locus Tag of target gene | Amplicon size (bp) |
|---|---|---|---|---|
| *qPCR* | | | | |
| adcAII (lmb) | adcAII-qPCR-F | aggttgatgtgtttgaagcg | M5005_Spy1711 | 106 |
| | adcAII-qPCR-R | gggtcataaagtgtcgcagg | | |
| cadD | cadD-qPCR-F | tctgatggagaagctattgcc | M5005_Spy1817 | 102 |
| | cadD-qPCR-R | attgtcagcaccacaacttgc | | |
| copA | copA-qPCR-F | tggtctcaggtcttgtggtc | M5005_Spy1405 | 97 |
| | copA-qPCR-R | gttgcagaaatggaggaagcc | | |
| holB | holB-qPCR-F | catagtaatcgtagtgggtttcgc | M5005_Spy1835 | 91 |
| | holB-qPCR-R | catttgaagtctgcttagaacgtg | | |
| siaA | siaA-qPCR-F | tgttgagggcatgtaccagtc | M5005_Spy1528 | 93 |
| | siaA-qPCR-R | tagtcctggtattgctggcg | | |
| tufA | tufA-qPCR-F | ggaccaatgccacaaactcg | M5005_Spy0508 | 109 |
| | tufA-qPCR-R | gcaactcttcgtcatcaacaagg | | |
| copY | copY-qPCR-F | gtcttatcggcaaggaagagc | M5005_Spy1406 | 108 |
| | copY-qPCR-R | tgttgccgctgacaaatacc | | |
| copZ | copZ-qPCR-F | gcaatcggtccaggtaaatttgg | M5005_Spy1404 | 92 |
| | copZ-qPCR-R | tggtatccttcaaagcacgc | | |
| copX | copX-qPCR-F | ctgacgttagcaggtgtgttcc | M5005_Spy1403 | 90 |
| | copX-qPCR-R | agtgcatcgtccttaaacacc | | |
| gapA (plr) | gapA-check-F | gtagttaaagttggtattaacgg | M5005_Spy0233 | 1008 |
| | gapA-check-R | tttagcaatttttgcgaagtactca | | |
| *Mutant construction* | | | | |
| copA | copA KO-F1 | ccggcctcgagccaaaaattcaatgcctacg | M5005_Spy1405 | 597 |
| | copA KO-R1 | taactataaactatttaaataacagattccctgc gattaaaaggaaaa | | |
| copA | copA KO-F2 | cgtatgtattcaaatatatcctcctcagcacaag aggcacaactca | M5005_Spy1405 | 674 |
| | copA KO-R2 | ccggcgaattctcgaataggatgtttcaagggta | | |
| kanR | kan-F | tatgttatacgccaactttg | - | |
| | kan-R | ctaaaacaattcatccagtaa | | |

**Supporting Table 4. List of strains and plasmids used in this study.**

| Bacterial strains | Description | Reference |
|---|---|---|
| *Escherichia coli* | | |
| MC1061 | *E. coli* laboratory cloning strain | (1) |
| *Streptococcus pyogenes* | | |
| 5448 | *S. pyogenes* invasive M1T1 strain | (2) |
| 5448Δ*copA* | *S. pyogenes* 5448 *copA*::*kan* deletion mutant | This study |
| 5448Δ*copA*$^{+}$ | *S. pyogenes* 5448 *copA*$^{+}$ marker rescue mutant | This study |

| Plasmids | Description | Reference |
|---|---|---|
| pHY304 | Temperature sensitive shuttle plasmid::Em$^{R}$ | (3) |
| pHY304-copAKO | pHY304 + *copA* knockout construct inserted at the XhoI and EcoRI site | This study |
