## Supporting Figures for "A role for glutathione in buffering excess intracellular copper in *Streptococcus pyogenes*"

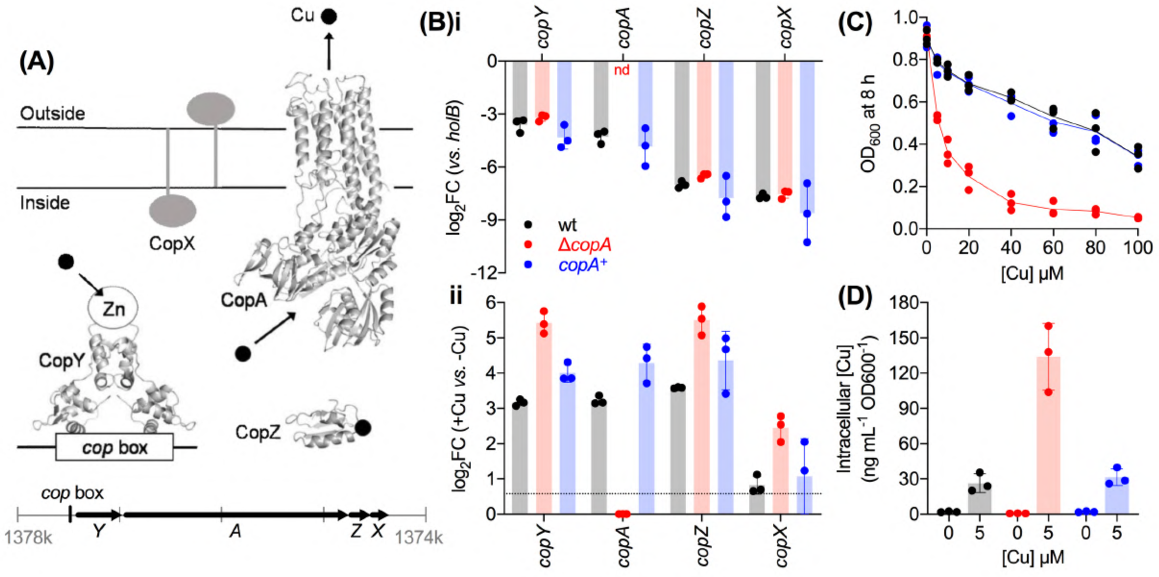


**Supporting Figure 1. Initial characterisation of GAS ∆*copA* mutant strain. (A) Working model of the Cop system*.*** Structural models were generated in SWISS-MODEL[1] using the following PDB IDs as templates: CopA (3j09), CopZ (1k0v), CopY (1fwz). Based on the *E. hirae* and *S. pneumoniae* paradigm, GAS CopY would bind Zn in its repressor form. Exchange of Zn with Cu during conditions of Cu excess would promote dissociation of CopY from its DNA binding site (*cop* box, ATTTACAAATGTAGTT)[2–4]. CopZ functions presumably as a cytoplasmic Cu-binding metallochaperone. CopA is a $P_{1B-1}$-type ATPase that extrudes excess Cu from the cytoplasm to the extracellular space. CopX has no known homologue outside GAS and has not been characterised. Organisation of the *copYAZX* operon is shown at the bottom*.* Gene map was generated using the genome sequence of GAS MGAS5005 (NCBI GenBank Accession No. CP000017.2, region *M5005_Spy_1406* to *M5005_Spy_1403*). **(B) (i) Basal and (ii) Cu-dependent transcription profiles of *cop* genes.** GAS was cultured with 0 µM (for panel (i); *n* = 3) or 5 µM (for panel (ii); *n* = 3) of added Cu. Bacteria were harvested at t = 4 h. Amounts of transcripts were quantified by qPCR and normalised to *holB* (for panel (i)) or to the corresponding untreated samples (for panel (ii)). nd, not detectable over background or non-specific amplification. **(C) Growth profiles.** GAS was cultured with added Cu as indicated (*n* = 3). Final $OD_{600}$ values were measured at t = 8 h. **(D) Levels of intracellular Cu.** GAS was cultured with added Cu as indicated (*n* = 3). Bacteria were harvested at t = 4 h and intracellular Cu levels were determined by ICP MS. All figures show GAS wild type (WT, black), ∆*copA* mutant (red), and *copA*⁺ complemented mutant (blue) strains.

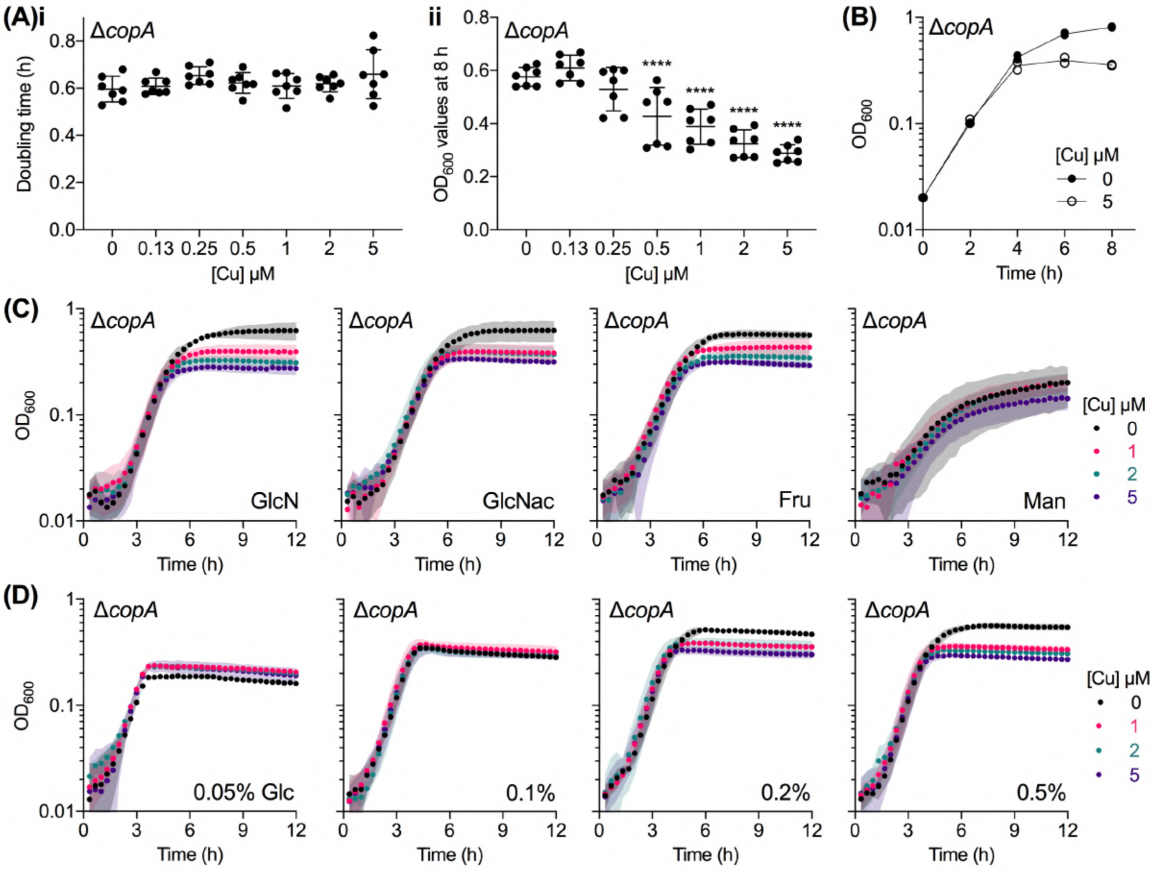


**Supporting Figure 2. Cu-dependent defects in growth. (A) Effects on (i) exponential doubling time and (ii) final culture density.** GAS $\Delta$*copA* mutant strain was cultured in microtitre plates with added Cu as indicated (*n* = 6). Doubling times were calculated by linear regression of $\log_2(OD_{600})$ values between t ~ 1 - 4 h. Final $OD_{600}$ values were determined at t = 8 h. $^{****}P < 0.0001$ (*vs.* untreated culture). **(B) Growth in bulk liquid cultures.** This experiment was performed to confirm that the growth defects observed in Figure 2A were not an artefact of bacterial culture in microtitre plates. GAS $\Delta$*copA* mutant strain was cultured in 40 mL of growth medium in 50 mL tubes, with added Cu salts as indicated (*n* = 2). $OD_{600}$ values were recorded every 2 h. **(C) Effect of carbon sources.** GAS $\Delta$*copA* mutant strain was cultured in microtitre plates with 0.5 w/v % each of glucosamine (GlcN), *N*-acetylglucosamine (GlcNac), fructose (Fru), or mannose (Man) as the sole carbon source, with added Cu as indicated (*n* = 3). **(D) Effect of glucose availability.** GAS $\Delta$*copA* mutant strain was cultured in microtitre plates with varying concentrations of glucose (w/v %) as the sole carbon source, with added Cu as indicated (*n* = 3).

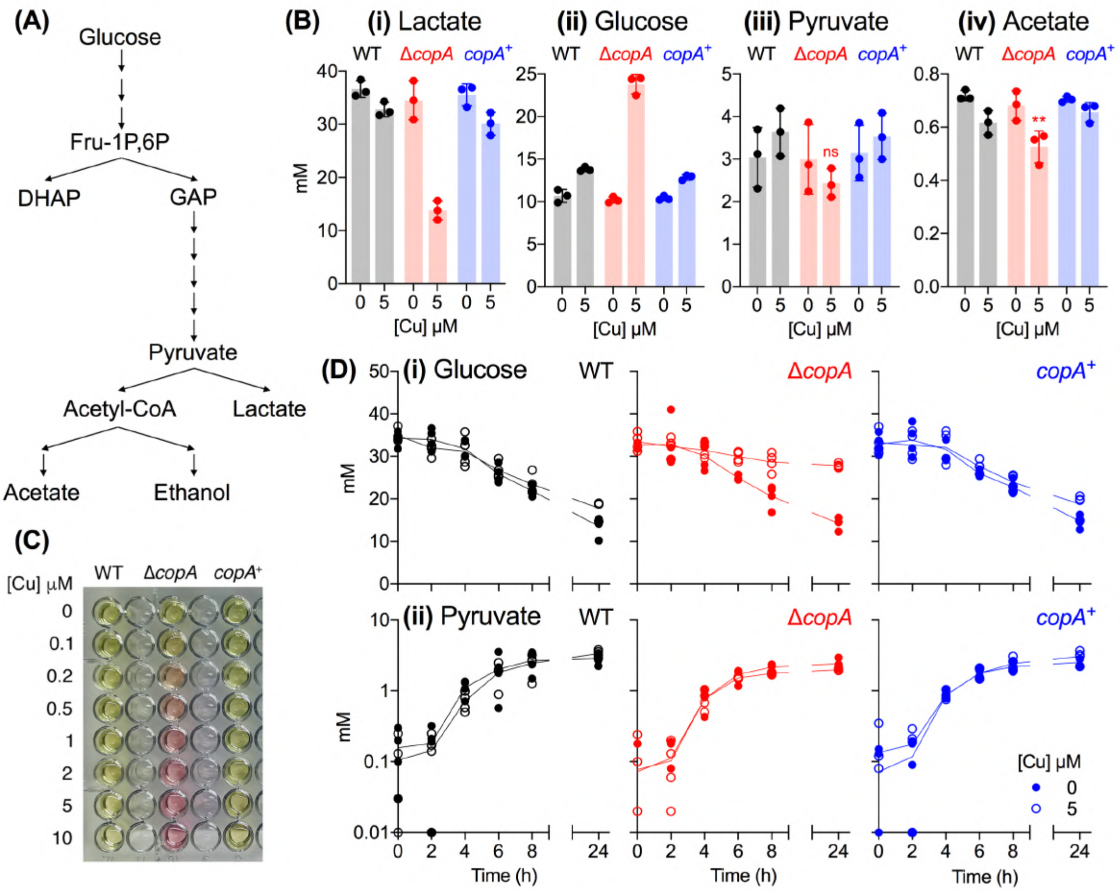


**Supporting Figure 3. Cu-dependent effects on lactic acid fermentation. (A) Fermentative pathways in GAS. (B) Levels of fermentation end products after 24 h of growth.** GAS was cultured with added Cu as indicated for t = 24 h ($n$ = 3). Amounts of **(i)** lactate, **(ii)** glucose, **(iii)** pyruvate, and **(iv)** acetate in the extracellular spent medium were determined. The results confirm that GAS carried out homolactic fermentation under our experimental conditions. Untreated cultures consumed 22.6 ± 0.5 mM of glucose and secreted 36 ± 2 mM of lactic acid (ca. 80% of the carbon yield), along with small amounts of pyruvate (3.1 ± 0.6 mM, ca. 7%) and acetate (0.7 ± 0.1 mM, ca. 1.5%). Cu treatment strongly suppressed lactate production ($P < 0.0001$) and glucose consumption ($P < 0.0001$). Cu treatment did not clearly affect pyruvate production ($^{ns}P = 0.64$) but it did have a small inhibitory effect on acetate production ($^{**}P = 0.002$). **(C) Media acidification.** Phenol red was present in our culture medium as a pH indicator. GAS was cultured with added Cu as indicated. After 24 h, the microtitre plates were centrifuged and the culture supernatants were transferred to a fresh microtitre plate and photographed. One representative photograph from numerous independent experiments is shown. **(D) Time-dependent production of fermentation end products.** GAS was cultured with added Cu as indicated ($n$ = 3). Cultures were sampled at the indicated time points and amounts of **(i)** glucose and **(ii)** pyruvate in the extracellular spent medium were determined. Cu treatment clearly affected glucose consumption in the Δ*copA* mutant ($P < 0.0001$).

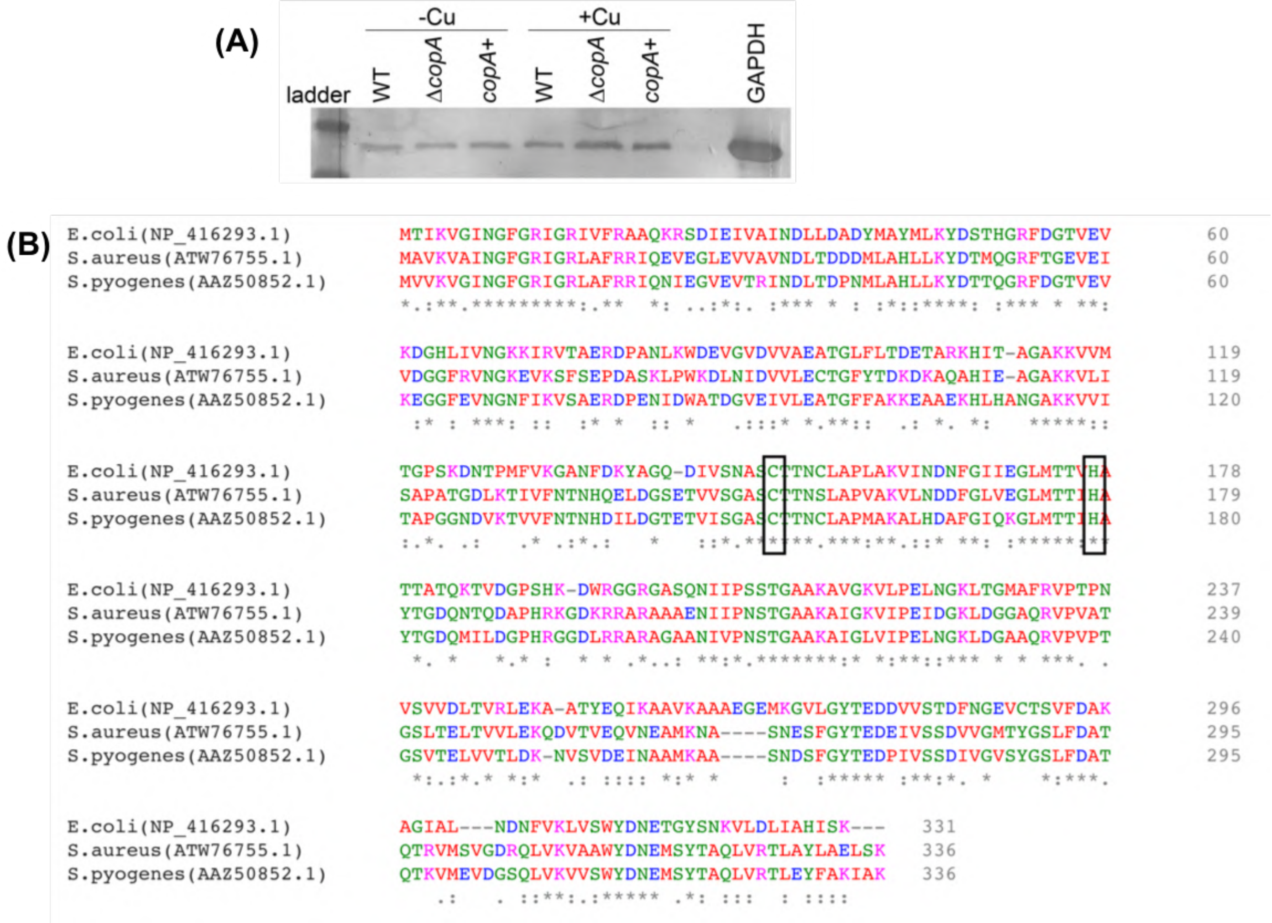


**Supporting Figure 4. (A) Western Blot analysis of GapA in cell extracts.** GAS was cultured without (-) or with (+) 5 µM of added Cu for t = 4 h. Bacteria were harvested and lysed by bead-beating. Exactly 5 µg of lysates were loaded and resolved on a 10% SDS PAGE gel using Tris-glycine running buffer. The gel was electroblotted onto a PVDF membrane. Rabbit α-GapA (1:1000; Walker AJ, Walker MJ, unpublished) was used for immunoblotting and detection of GapA (GAPDH). Rabbit α-IgG-HRP conjugate (1:1000, Sigma) was used as secondary antibody. The membrane was developed with 3,3'-diaminobenzidine and hydrogen peroxide. Purified recombinant GapA (Walker AJ, Walker MJ, unpublished) was used as a positive control. A representative blot from three independent experiments is shown. **(B) Conservation of amino acid sequences between GapA homologues from *E. coli*, *S. aureus*, and GAS.** The Cys active site and the conserved His metal-binding ligand are boxed for clarity. NCBI GenBank accession numbers are shown in brackets.

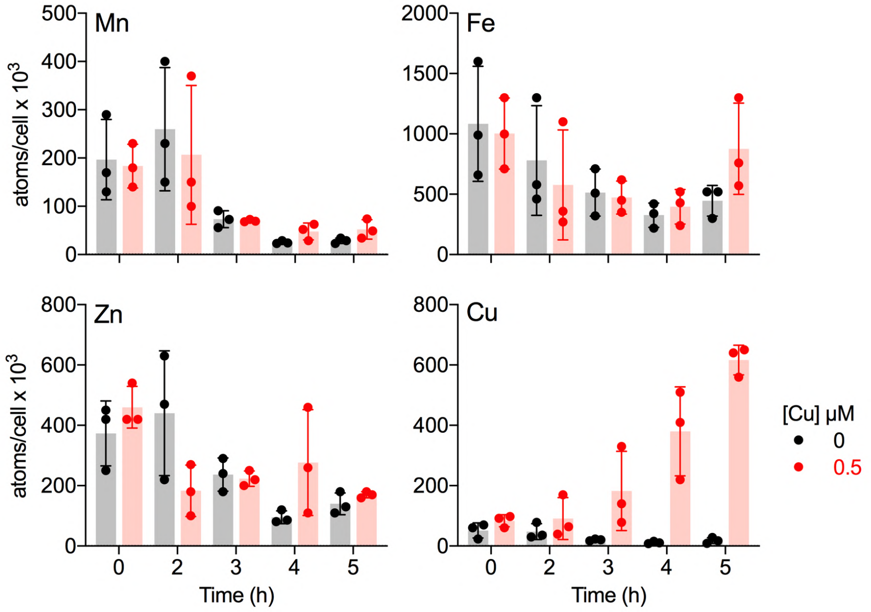


**Supporting Figure 5. Time-dependent changes in intracellular Cu levels.** GAS Δ*copA* mutant strain was cultured with 0 μM (black) or 0.5 μM (red) of added Cu (*n* = 3). Cultures were sampled at the indicated time points, plated out for colony counting, and subjected to metal analyses by ICP MS. Total levels of intracellular Mn, Fe, Zn, and Cu are presented as number of metal atoms per cell.

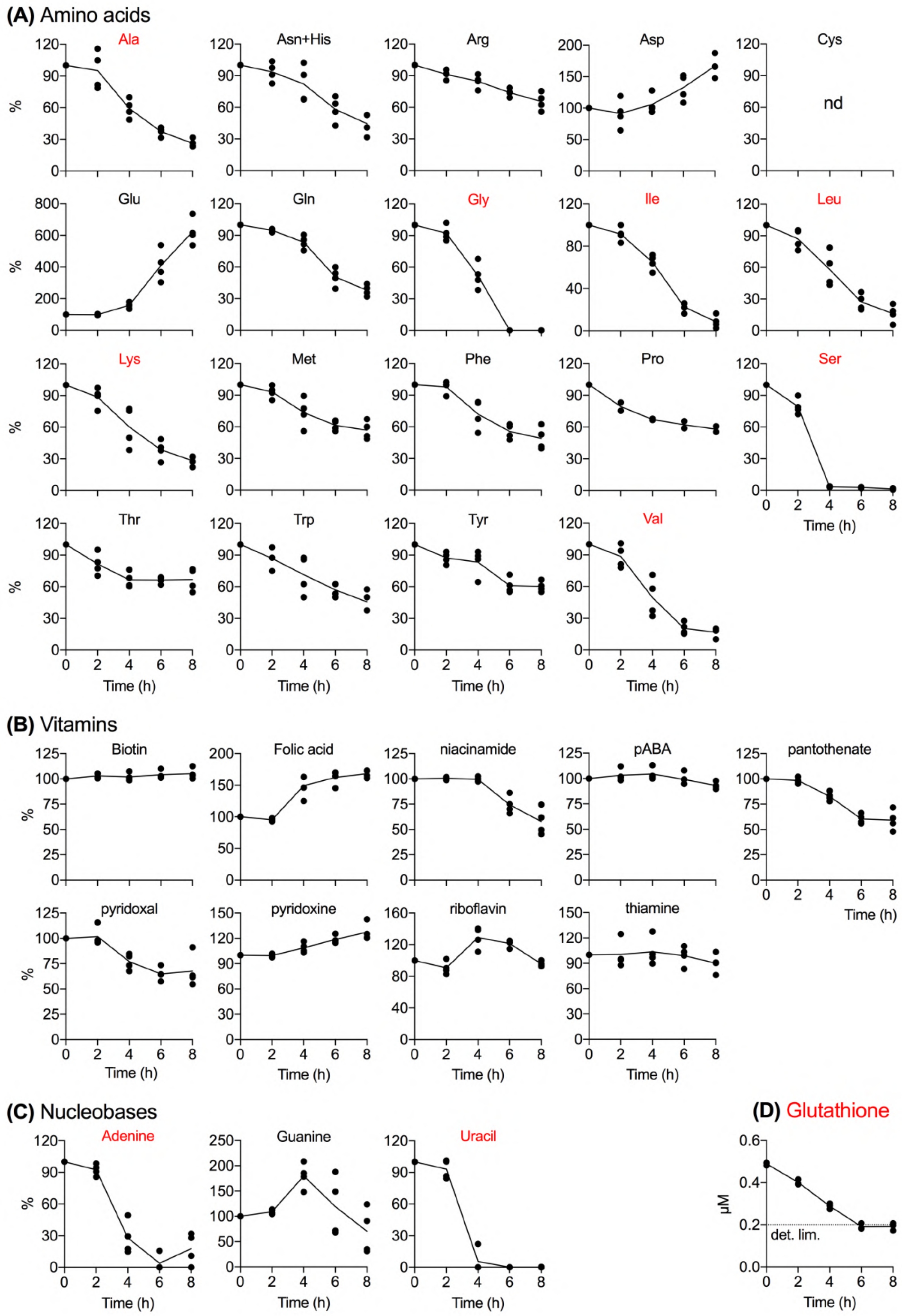
(A) Amino acids
Ala
Asn+His
Arg
Asp
Cys
nd
Glu
Gln
Gly
Ile
Leu
Lys
Met
Phe
Pro
Ser
Thr
Trp
Tyr
Val
%
Time (h)
(B) Vitamins
Biotin
Folic acid
niacinamide
pABA
pantothenate
pyridoxal
pyridoxine
riboflavin
thiamine
(C) Nucleobases
Adenine
Guanine
Uracil
(D) Glutathione
µM
det. lim.

**Supporting Figure 6. Time-dependent changes in the levels of extracellular growth medium components: (A) amino acids; (B) vitamins; (C) nucleobases; and (D) glutathione.** GAS Δ*copA* mutant strain was cultured without added Cu ($n$ = 4). The cultures were centrifuged (5000 x $g$, 10 min) and filtered through a 0.22 μm PES membrane at the indicated time points. The filtrates (spent media) were subjected to analyses by LC-MS on a Sciex QTRAP 6500 at Durham Biosciences Proteomics Facility. The amount of each amino acid, vitamin, and nucleobase in these filtrates was normalised to the amounts present at t = 0 h. The amino acid Cys was below detection limit. Glutathione levels were measured separately by Gor/DTNB recycling assay[5]. Medium components that were nearly or completely spent at t = 4 h were highlighted in red.

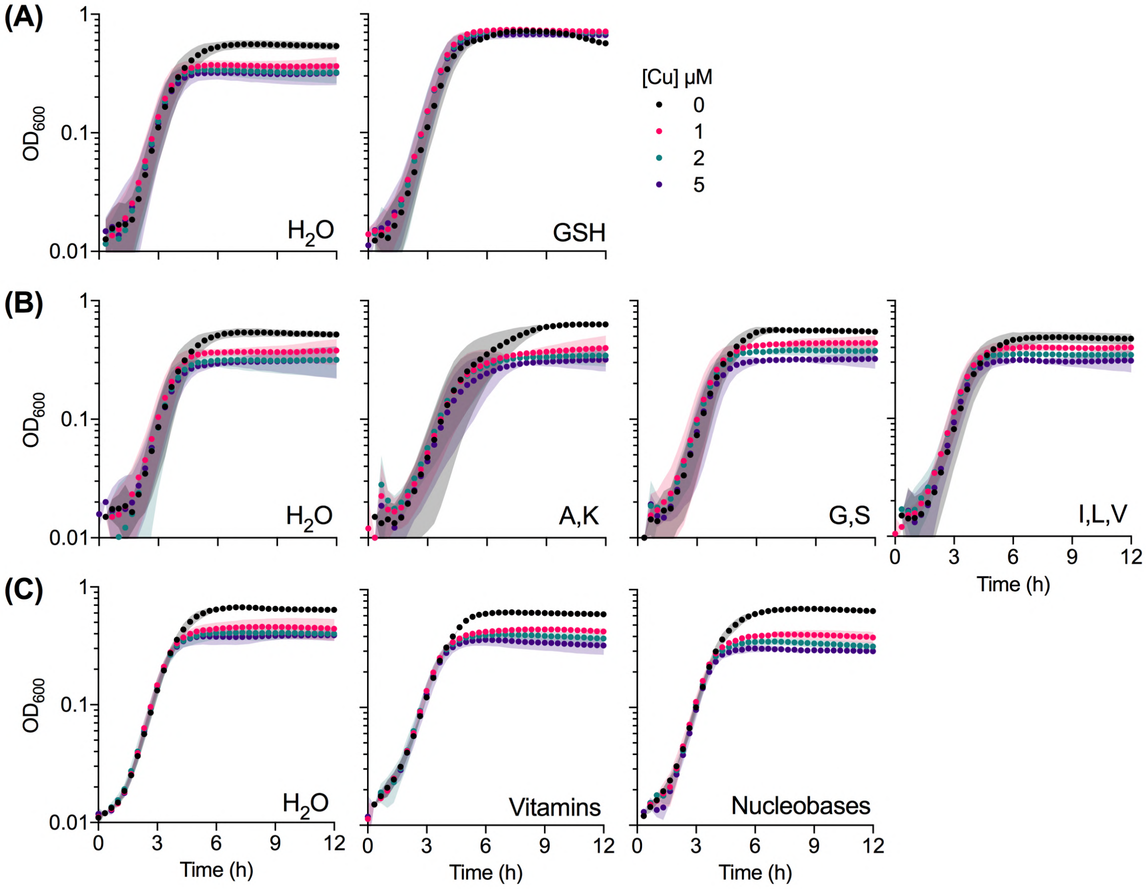


**Supporting Figure 7. Effects of media supplements on bacterial growth.** GAS Δ*copA* mutant strain was cultured with added Cu as indicated. The cultures were also supplemented with: **(A) GSH** (1 mM, *n* = 3); **(B) combinations of select amino acids** (1 mM each, *n* = 3, single-letter amino acid codes are shown); or **(C) vitamins and nucleobases** (to 5X the initial concentration each, *n* = 3). The corresponding untreated (water) control cultures for each experiment are shown on the left.

```
S.mutans(WP_002262138.1)      MKLKHILRIGAVAFASILLLTACGSKT--------SKKTV       32
S.pyogenes(AAZ50889.1)        MIKKQYLGRATIALASTLVLAACGSSKTAESGNQGSSKEV       40

S.mutans(WP_002262138.1)      TLATVGTTNPFSYEKKGKLTGYDIEVAKEVFKASDKYDVK       72
S.pyogenes(AAZ50889.1)        LFATVGTTAPFSYEKGGQLTGYDIEVAKAVFKGSDDYKVS       80

S.mutans(WP_002262138.1)      YQKTEWTSIFSGLDSDKYQIGANNISYTKERANKYLYSNP       112
S.pyogenes(AAZ50889.1)        FKKTEWSSIFTGLDSGKYQMGGNNISFTKERSAKYLFSYP       120

S.mutans(WP_002262138.1)      TASNPLVLVVPKDSDIKSYNDIAGHSTQVVQGNTTVSMLQ       152
S.pyogenes(AAZ50889.1)        IGSTPSVLVVPKDSDIKSFDDIQGHTTQVVQGTTSVAQLE       160

S.mutans(WP_002262138.1)      KFNKNHENNQVKLNFTSEDLAHQIRNVSDGKYDFKIFEKI       192
S.pyogenes(AAZ50889.1)        DFNKKHSDNPVTLKFTNENITQMLTNLSEGKADFKIFDAP       200

S.mutans(WP_002262138.1)      SAETIIKEQGLDNLKVIDLPSDQKPYVYFIFAQDQKDLQK       232
S.pyogenes(AAZ50889.1)        TVNAIIKNQGLDNLKTIELTSTEQPFIYFIFSQDQEKLQS       240

S.mutans(WP_002262138.1)      FVNKRLKKLYENGTLEKLSKKYLGGSYLPDKKDMK-----       267
S.pyogenes(AAZ50889.1)        FVNKRIKELTADGTLSKLAKEHLGGDYVPSDKELKLPTAN       280
```

**Supporting Figure 8. Alignment of GshT protein sequences from *S. mutans*[6] and GAS.** The two proteins share 59% sequence identity (in red) and overall 74% sequence similarity (in blue). NCBI GenBank accession numbers are shown in brackets.

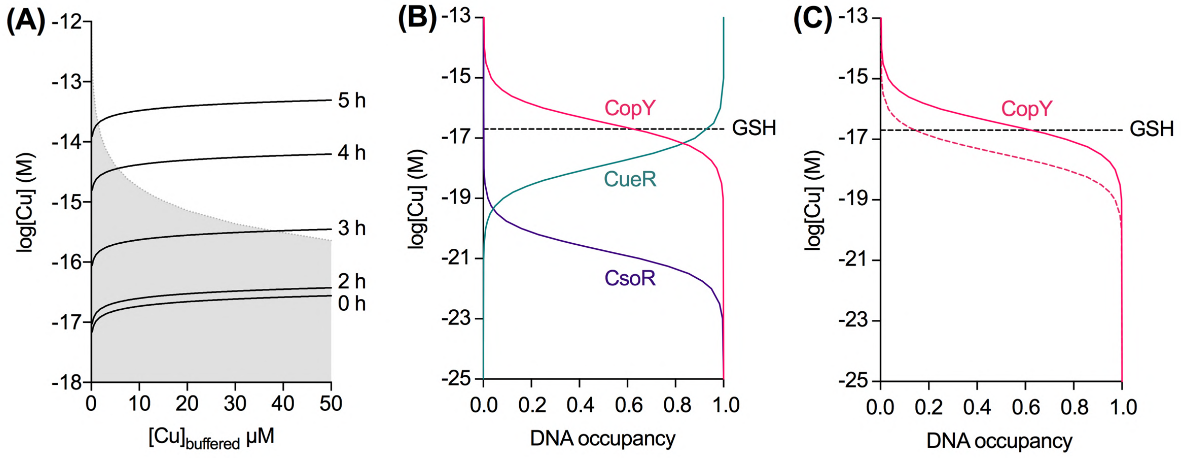


**Supporting Figure 9. Buffering of intracellular Cu by GSH. (A) Time-dependent and GSH-dependent changes in the ability of GSH to buffer intracellular Cu.** The threshold of Cu availability imposed by GSH (plotted as log[Cu]) was estimated using equation 4 in Morgan *et. al.* (2017)[7]. Intracellular pH was assumed to be 7.2. Intracellular GSH concentrations at each time point were measured in Figure 6A ([GSH] = 4.1, 3.3, 0.76, 0.11, 0.03 mM at t = 0, 2, 3, 4, 5 h, respectively). This estimate is valid only in the region shaded in grey, i.e. where GSH is present in a huge excess ([GSH]/[Cu] >> 20)[7]. **(B) Comparison of GSH with characterised bacterial Cu sensors.** The thresholds of Cu availability imposed by CsoR from *B. subtilis*[8], CopY from *S. pneumoniae*[2], and CueR from *S. Typhimurium*[9] were estimated using the calculator developed by Osman *et al.*[9] (Supporting Table 5). The midpoint threshold imposed by 4 mM GSH is shown as straight dashed line. **(C) Effect of Cu affinity on the CopY sensor.** The thresholds of Cu availability imposed by CopY were estimated using the using published Cu binding affinity of log $K_{Cu}$ = 16.6 (solid pink curve)[2] and a 10X tighter affinity of log $K_{Cu}$ = 17.6 (dashed pink curve). The midpoint threshold imposed by 4 mM GSH is shown as straight dashed line.

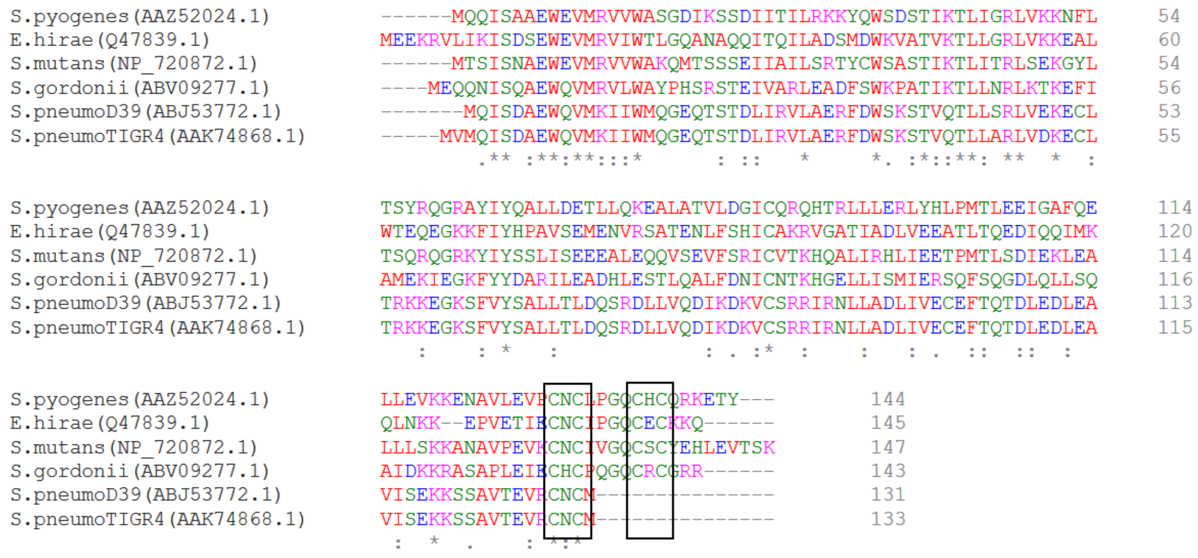


**Supporting Figure 10. Alignment of CopY protein sequences.** The two Cys-X-Cys motifs are boxed for clarity. NCBI GenBank accession numbers are shown in brackets.

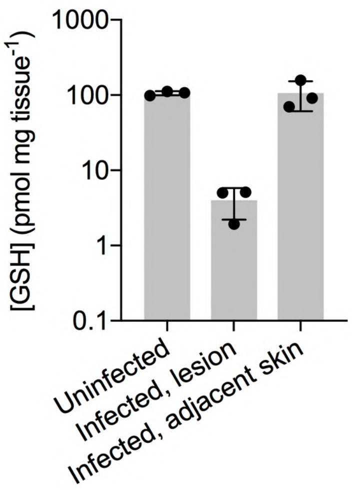


**Supporting Figure 11. Amounts of host GSH in the ulcerative lesions.** Mice were infected subcutaneously with GAS wild type strain. After 3 days, the skin of uninfected mice (naive), skin lesions of infected mice, and control skin adjacent to the lesions of infected mice were excised (*n* = 3 each). Total GSH levels were measured and normalised to the weight of the tissue sample. There was a clear reduction in GSH levels in the lesions from infected mice when compared with healthy skin from uninfected mice (*P* = 0.0059) and adjacent healthy skin from infected mice (*P* = 0.0059).
